# One Thousand SARS-CoV-2 Antibody Structures Reveal Convergent Binding and Near-Universal Immune Escape

**DOI:** 10.1101/2025.08.07.669152

**Authors:** Zirui Feng, Zhe Sang, Yufei Xiang, Alba Escalera, Adi Weshler, Dina Schneidman-Duhovny, Adolfo García-Sastre, Yi Shi

**Affiliations:** Center of Protein Engineering and Therapeutics, Department of Pharmacological Sciences, Icahn School of Medicine at Mount Sinai, New York, NY, USA, 10029; Department of Microbiology, Icahn School of Medicine at Mount Sinai, New York, NY, USA, 10029; Global Health and Emerging Pathogens Institute, Icahn School of Medicine at Mount Sinai, New York, NY, USA, 10029; School of Computer Science and Engineering, The Hebrew University of Jerusalem, Jerusalem, Israel; Department of Medicine, Division of Infectious Diseases, Icahn School of Medicine at Mount Sinai, New York, NY, USA, 10029; The Tisch Cancer Institute, Icahn School of Medicine at Mount Sinai, New York, NY, USA, 10029; Department of Pathology, Molecular and Cell-Based Medicine, Icahn School of Medicine at Mount Sinai, New York, NY 10029, USA; The Icahn Genomics Institute, Icahn School of Medicine at Mount Sinai, New York, NY, USA, 10029

**Author notes:** Contribute equally (Zirui Feng, Zhe Sang). Contact: Yi Shi.

## Abstract

Since the emergence of SARS-CoV-2, understanding how antibodies recognize and adapt to viral evolution has been central to vaccine and therapeutic developments. To date, over 1,100 SARS-CoV-2 antibody structures—16% of all known antibody–antigen complexes—have been resolved, marking the largest structural biology effort towards a single pathogen. Here, we present a comprehensive analysis of this landmark dataset to investigate the principles of antibody recognition and immune escape. Human immunoglobulins (IgGs) and camelid single-chain antibodies dominate the dataset, collectively mapping 99% of the receptor-binding domain surface. Despite remarkable sequence and conformational diversity, antibodies exhibit striking convergence in their paratope structures, revealing evolutionary constraints in epitope selection. Structural and functional analyses reveal near-universal immune escape of antibodies, including all clinical monoclonals, by advanced variants such as KP3.1.1. On average, over one-third of antibody epitope residues are mutated. These findings support pervasive immune escape, underscoring the need to effectively leverage multi-epitope targeting strategies to achieve durable immunity.

## Main Text

Five years after its emergence, SARS-CoV-2 has claimed millions of lives and spurred an unprecedented global effort to develop countermeasures, including vaccines, small-molecule inhibitors, and therapeutic antibodies (*1, 2*). Structural biology has played a central role in this response, revealing key molecular features of viral infection, evolution, and immune recognition, with insights that have directly informed therapeutic and vaccine design (*3*). As part of this effort, high-resolution structures of numerous neutralizing antibodies, including 9 approved by the U.S. Food and Drug Administration (FDA), and 1 by the European Medicines Agency (EMA), and 3 in East Asia—have been rapidly resolved in complex with viral antigens (*4–15*). These antibodies are among the most potent antiviral agents developed to date, offering advantages over small molecules such as high specificity, extended half-lives, and the ability to engage host immune effector functions for viral clearance(*16, 17*).

Yet the virus continues to evolve, accumulating mutations throughout its genome. Many of these changes affect the spike glycoprotein, particularly the receptor-binding domain (RBD), a critical site for viral entry and the primary target of neutralizing antibodies (*18, 19*). Although the detailed mechanisms driving this evolution remain incompletely understood, these mutations have enabled significant immune escape, undermining the efficacy of both vaccines and antibody therapies (*20–23*). New variants have emerged every 3 to 6 months, frequently outpacing the development and deployment of updated countermeasures, highlighting the limitations of reactive, update-based strategies (*20, 24*). The ongoing risk of zoonotic spillover from related coronaviruses further emphasizes the need for durable, broadly protective antiviral therapeutics (*25, 26*).

The growing collection of SARS-CoV-2 antibody structures, though still not fully explored, offers a unique opportunity to examine how antibody recognition evolves under selective pressure(*6, 13, 27*). Here, we present a comprehensive analysis of this massive dataset, focusing on two dominant antibody classes: human IgGs and camelid single-chain V_H_Hs (nanobodies/Nbs)(*28–32*). Through systematic structural comparisons, we identify distinct modes of epitope targeting and shared molecular principles that underpin high-affinity binding. Despite striking sequence diversity, antibodies consistently converge on a limited set of structural solutions for epitope recognition, reflecting evolutionary and biophysical constraints. We further show that immune escape is near-universal and provide structural insights into why numerous antibodies including all clinical ones and antibodies targeting conserved epitopes may fail. These findings define key principles of antiviral antibody recognition and offer a roadmap for designing next-generation therapeutics with improved durability.

### Rapid deposition of 1,100+ SARS-CoV-2 antibody structures

As of June 2024, the SAbDab database includes 6,892 resolved antibody–antigen complex structures(*33*), with 41% targeting viral antigens and 39% against mammalian proteins, highlighting both immunological relevance and biomedical applications (**Fig. 1A**). Nearly all structures were determined by X-ray crystallography or cryo-electron microscopy (cryo-EM), with median resolutions of 2.5 Å and 3.3 Å, respectively (**fig. S1A, S1B**).

**Fig. 1:**
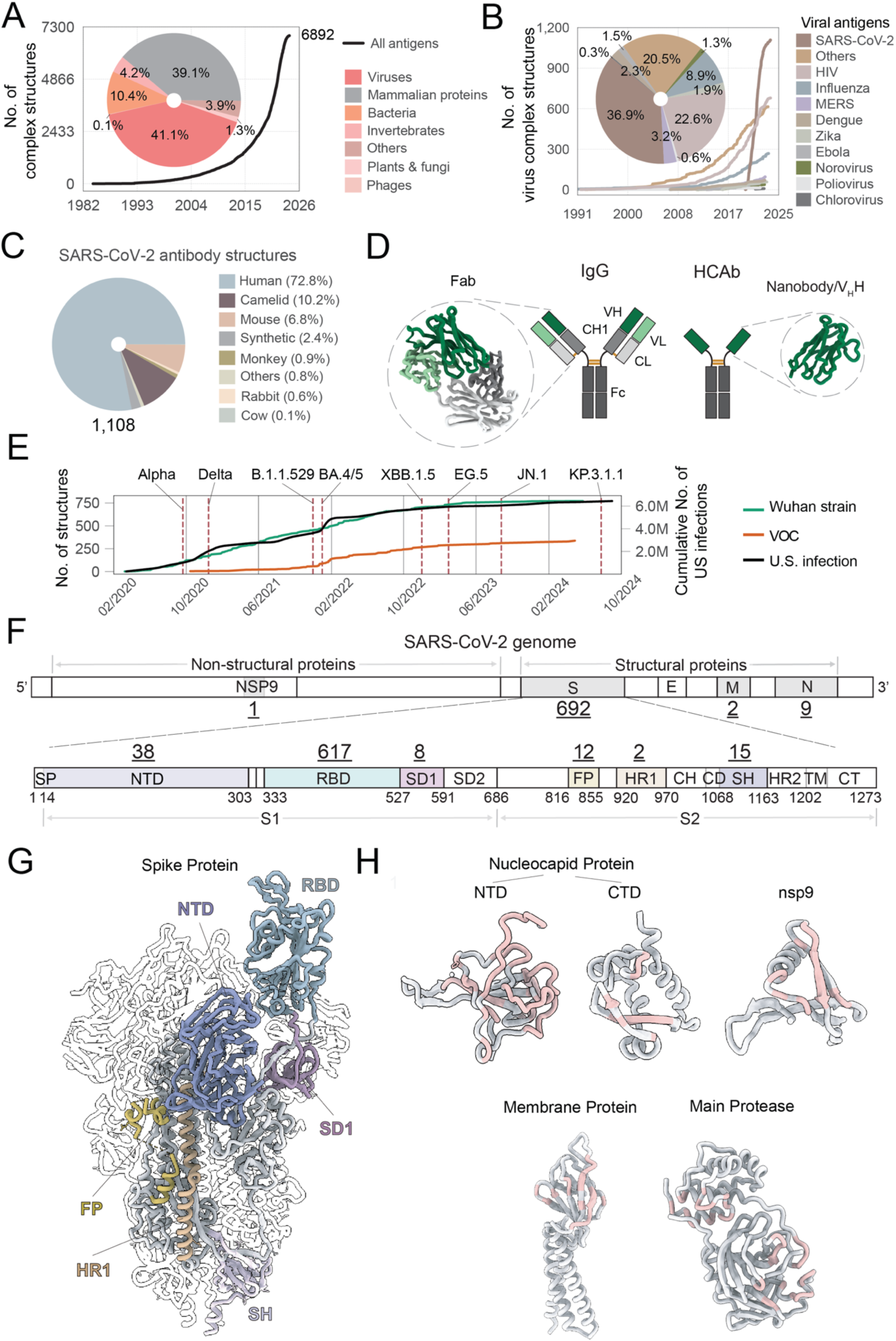
Overview of SARS-CoV-2 antibody PDB structures. (A) Cumulative count of antibody-antigen complex structures deposited in PDB. The inset pie chart shows distribution of structures by taxonomic groups of antigens. (B) Cumulative count of antibody antigen complex structures for viral proteins. The inset pie chart shows distribution of structures by virus species. (C) Pie chart showing species of origin for structurally determined antibodies in complex with SARS-CoV-2 functional proteins. (D) Schematics of monoclonal IgG and camelid heavy chain antibody (HcAb). Zoom-in view shows representative protein structures of the Fab domain (PDB: 7KMG) from IgG and the Nb domain (PDB: 7N9A) from HcAb. (E) Cumulative count of antibody structures determined in complex with either SARS-CoV-2 wildtype or variant, compared with the scaled cumulative count of infections in the U.S. since the pandemic. The emergence of several variants of concerns is labeled. (F) Schematic representation of the SARS-CoV-2 genome and the corresponding number of antibodies targeting each functional protein. In the genome schematic, S denotes the spike protein, E the envelope protein, M the membrane protein, and N the nucleocapsid protein. A zoom-in view shows different domains on the spike protein, SP: Signal peptide, NTD: N-terminal domain, RBD: Receptor binding domain, SD1: S1 subdomain 1, SD2: S1 subdomain 2, FP: Fusion peptide, HR1: Heptad repeat 1, CH: Central helix, CD: Connector domain, SH: S2 stem helix, HR2: Heptad repeat 2, TM: Transmembrane, CT: Cytoplasmic tail. (G) Cartoon representation of SARS-CoV-2 spike proteins (PDB: 6X2A). Domains targeted by antibodies are colored, including NTD, RBD, SD1, FP, HR1 and SH. (H) Cartoon representations of antibody-targeted SARS-CoV-2 proteins, including the nucleocapsid protein N-terminal domain (NTD; PDB: 7CR5) and C-terminal domain (CTD; PDB: 7NOI), non-structural protein 9 (nsp9; PDB: 8DQU), membrane protein (PDB: 7VGR), and main protease (PDB: 6LU7). Antibody epitopes are colored in pink.

Among more than 700 antigen targets, SARS-CoV-2 has become the most extensively characterized, with 1,108 structures corresponding to 705 unique antibodies, representing 36.9% of all viral antigen structures. This surpasses datasets for HIV (22.6%), influenza (8.9%), MERS (3.2%), and dengue (2.3%) (**Fig. 1B, fig. S1C, S1D**). The deposition rate closely tracked the pandemic’s course, peaking in early 2023, approximately one year after Omicron emerged (**Fig. 1E**).

Human IgGs comprise 72.8% of the dataset, while Nbs account for 10.2% (**Fig. 1C, 1D**), reflecting increased interest in Nbs for their small size, stability, and amenability to engineering(*34*). The majority of SARS-CoV-2 antibodies target the spike glycoprotein (692 antibodies), especially the RBD (617 antibodies; **fig. S1E, S1F**), with fewer directed at the N-terminal domain (38), stem helix (15), and fusion peptide (12) (**Fig. 1F, 1G**). By contrast, only a handful target other structural proteins such as the nucleocapsid (9) and membrane protein (2), or non-structural proteins such as Nsp9 and Mpro (1) (**Fig. 1H, fig. S1G**). The remarkable structural diversity of RBD antibodies offers an unparalleled resource to study molecular architecture of antibody binding and the mechanisms underlying viral escape.

### 99% of RBD surface residue is antigenic

To enable systematic, large-scale analysis of SARS-CoV-2 antibody recognition, we classified all RBD-targeting IgGs and Nbs into four broadly defined epitope groups based on residue overlap (**Fig. 2A–C, Methods**) (*35*). Groups I and II target the highly variable receptor-binding site (RBS)(*19*), while Groups III and IV engage more conserved, non-RBS regions. Epitope usage between IgGs and Nbs was highly correlated in Groups I, II, and IV (Pearson r > 0.8), whereas Group III showed minimal overlap: IgGs predominantly bound residues 333–348, 440–462, and 478–493, while Nbs preferentially targeted 380–394, 426–430, and 514–525 (**fig. S2A–D**).

**Fig. 2:**
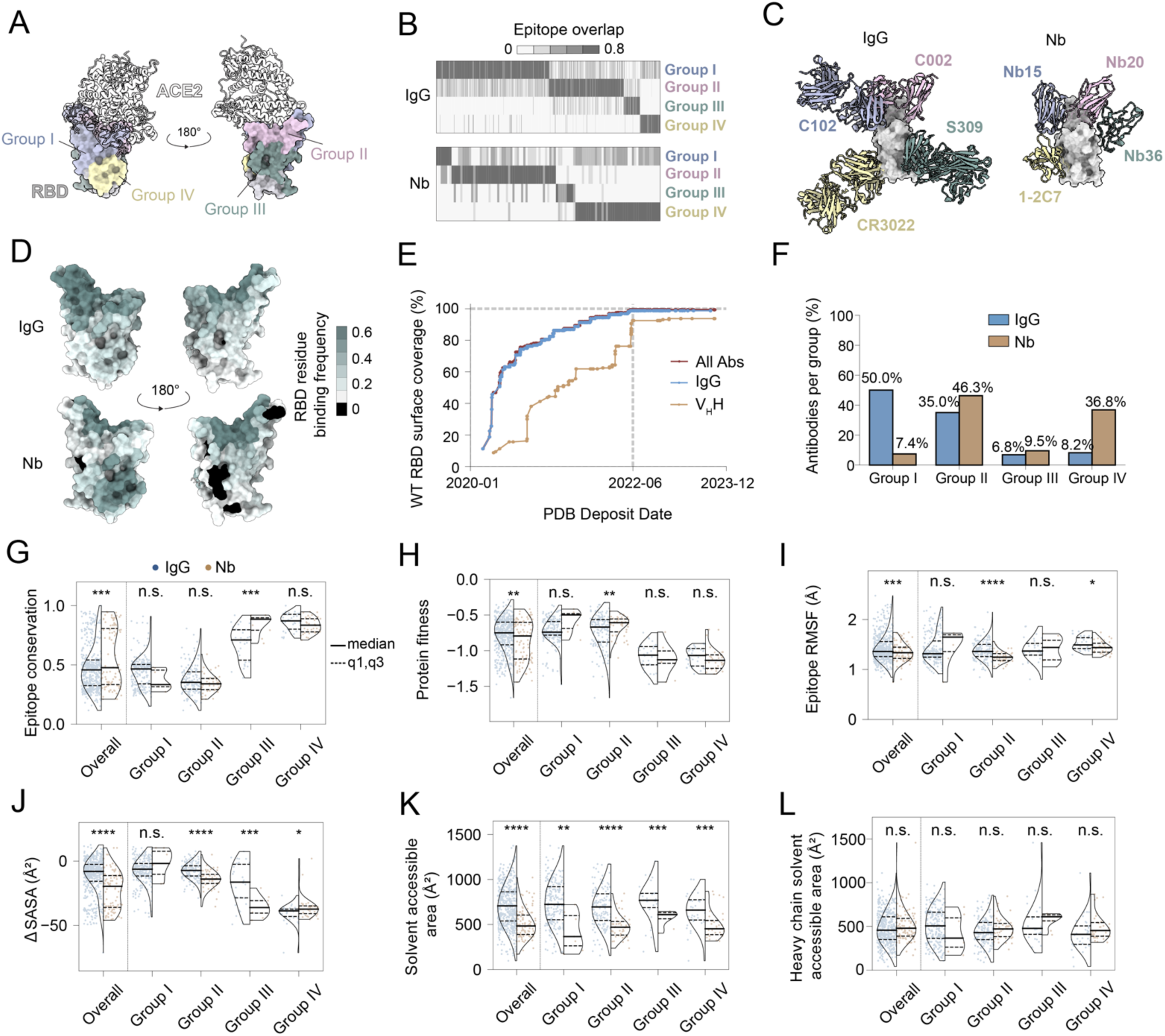
Summary of antibody epitope classification, coverage and properties on SARS-CoV-2 RBD: (A) Summary of four epitope groups (**Methods**) on RBD targeted by antibodies (PDB:6M0J). Human receptor ACE2 is shown in cartoon representation. (B) Heatmaps showing similarity between antibody epitopes and 4 reference epitope groups shown in (A). Upper panel: IgG, lower panel: Nb. (C) Superposition of representative IgGs and Nbs structures (Cartoon representation) in complex with RBD (surface representation). Class 1 (blue): C102 (PDB: 7K8M) and Nb15 (PDB: 8H5T); Class 2 (pink): C002 (PDB: 7K8T) and Nb20 (PDB: 7JVB); Class 3 (green): S309 (PDB: 8HWS) and Nb36 (PDB: 7MEJ); Class 4 (yellow): CR3022 (PDB: 6YLA) and 1-2C7 (PDB: 7X2M). (D) Surface representation of RBD (PDB:6M0J), the RBD residues are colored based on binding frequency targeted by IgG (upper) or Nb (lower). (E) Cumulative plot showing WT RBD surface coverage by antibodies over time. (F) Bar plot showing distribution of IgGs and Nbs across four groups. (G) Comparative analysis of epitope sequence conservation. Violin plots showing epitope sequence conservation and the pairwise statistical significance by epitope group. The bold line represents the median, and the dashed lines represent the 25th, 75th percentile. IgG: blue; Nb: orange. Statistical analysis was performed using a two-tailed student t test, \**p* < 0.05, \*\**p* < 0.01, \*\*\**p* < 0.001, \*\*\*\**p* < 0.0001. (H) Comparative analysis of epitope protein fitness. Violin plots showing epitope protein fitness and the pairwise statistical significance by epitope group. (I) Comparative analysis of epitope structural flexibility. Violin plots showing epitope flexibility (RMSF by MD simulation) and the pairwise statistical significance by epitope group. (J) Comparative analysis of epitope accessibility in context with RBD dynamics on Spike. Violin plots showing epitope accessibility change from up to down conformation and the pairwise statistical significance by epitope group. (K) Comparative analysis of epitope solvent accessible area. Violin plots showing epitope solvent accessible surface area and the pairwise statistical significance by epitope group. (L) Comparative analysis of heavy chain epitope solvent accessible area. Violin plots showing heavy chain epitope solvent accessible surface area and the pairwise statistical significance by epitope group.

Striking differences emerged in epitope distribution. Half of all IgGs were RBS-directed (Group I), compared to just 7.4% of Nbs. In contrast, Group IV Nbs, which target conserved, non-RBS regions, were 4.5 times more frequent than their IgG counterparts (36.8% vs. 8.2%) (**Fig. 2F**). Collectively, IgGs and Nbs cover 99% and 94% of the RBD surface, respectively (**Fig. 2D–E, fig. S2E**). Although antibodies are known to recognize specific antigenic motifs (*36*), our study provides the first structural evidence that virtually every solvent-accessible residue on a viral domain can be targeted by mammalian antibodies. Saturation of the RBD antigenic landscape highlights the remarkable flexibility of the humoral immune response and offers a foundation for identifying conserved and potentially synergistic epitopes. Achieving this comprehensive structural mapping within just two years of the pandemic outbreak underscores the remarkably rapid scientific response.

### Nbs prefer conserved and cryptic epitopes through distinct binding strategies

To probe the molecular basis of epitope selection, we assessed conservation, mutational fitness, and structural dynamics(*37, 38*) (**Table S2**). Conservation was quantified using amino acid variability across 19 sarbecovirus RBD sequences (*39*) (**fig. S3A**), revealing that Nbs are significantly enriched for conserved epitopes (**Fig. 2G**), primarily due to their overrepresentation in Groups III and IV (**Fig. 2F**). Nbs also preferentially target regions of high viral fitness, which tolerates fewer mutations (**Fig. 2H, fig. S3B**). As expected, conservation and mutational tolerance were inversely correlated (Pearson r = –0.85, **table S1**).

Molecular dynamics simulations showed that Nbs bind more rigid regions, with lower average RMSF values compared to IgGs (**Fig. 2I, fig. S3C**), and their epitopes undergo larger accessibility changes between the “up” and “down” RBD conformations(*40*), indicating preferential recognition of cryptic, conformationally gated surfaces (**Fig. 2J, fig. S3D**) (*41, 42*). The median surface area of Nb epitopes (479 Å²) was significantly smaller than that of IgGs (714 Å²) (**Fig. 2K)** and resembled the footprint of IgG heavy chains alone (**Fig. 2L**).

Mechanistically, Nbs exploit a distinct binding mode to access these conserved and cryptic epitopes. Unlike IgGs, which rely predominantly on aromatic contacts via tyrosine, Nbs also engage targets using electrostatic interactions, including arginine-mediated salt bridges and cation–π contacts (**Supplemental information for detailed analysis, figs. S4, S5**). These adaptations likely compensate for the absence of a light chain and enable compact paratopes to achieve high-affinity binding(*43–48*). Together, these findings highlight how physiochemical properties shape differential epitope targeting between IgGs and Nbs, with implications for antibody durability and rational engineering.

### Structural convergence amid CDR3 diversity enables epitope selection

Among antibody CDR loops, the heavy chain CDR3 is the most diverse in sequence and structure and plays a dominant role in antigen recognition (*48*). Superposition of antibody–RBD complexes revealed extensive CDR3 conformational variability in both orientation and geometry across all four epitope groups and for both IgGs and Nbs (**Fig. 3A–B**). To investigate how structurally diverse antibodies recognize shared epitopes, we clustered antibodies within each group into sub-clusters with at least 40% epitope residue overlap (**Methods**). This yielded 34, 33, 11, and 7 sub-clusters for IgGs across Groups I–IV, and 5, 10, 5, and 8 for Nbs, respectively (**Fig. 3C–D**).

**Fig. 3:**
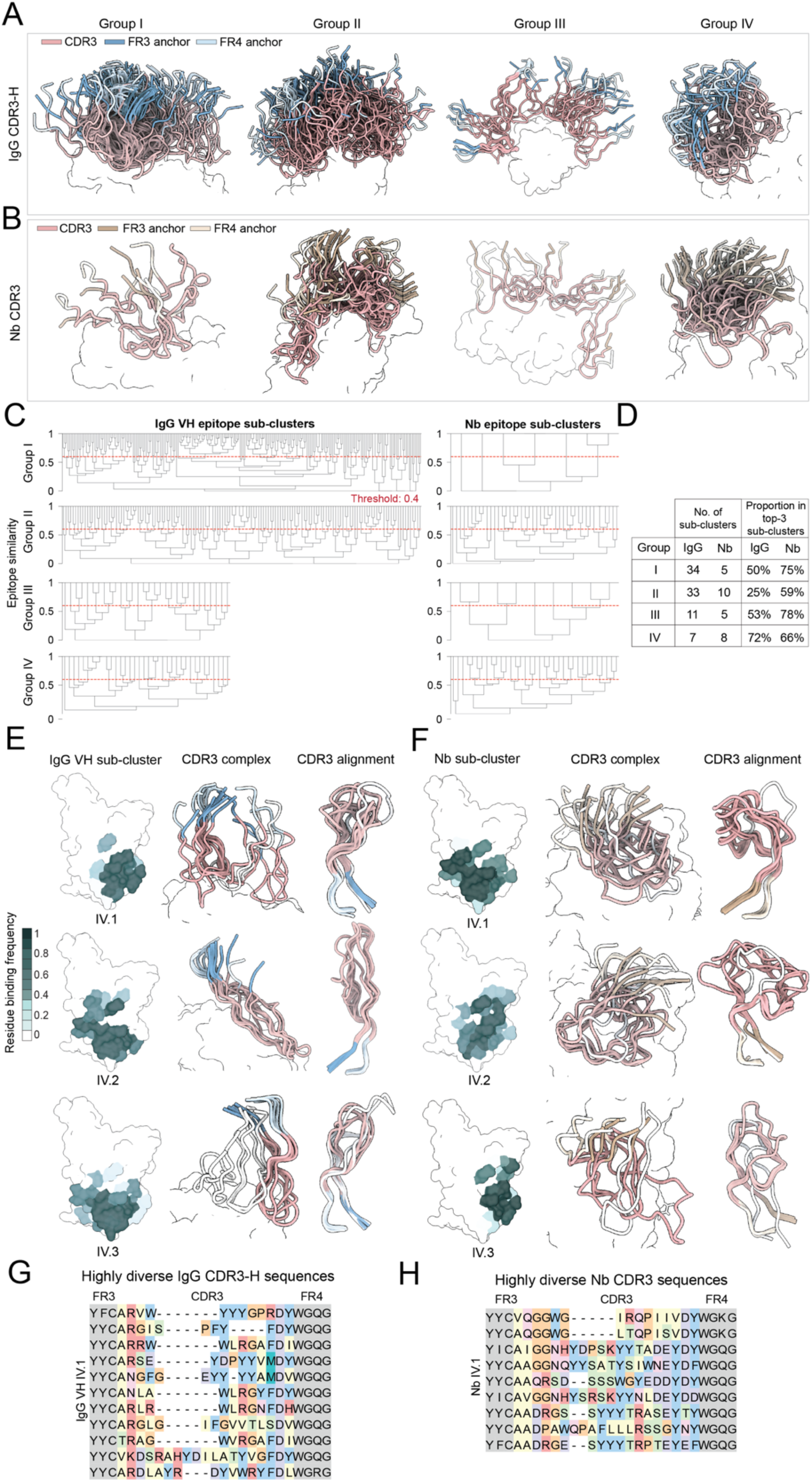
Epitope sub-clusters and CDR3 loop conformation convergence. (A) Superpositions of IgG CDR3-H loops (Cartoon representation) in complex with the RBD (surface representation) for each epitope group defined in Fig.2. CDR3 regions are colored in pink and FR3/FR4 anchors are colored in blue and light blue. (B) Superpositions of Nb CDR3 loops (Cartoon representation) in complex with the RBD (surface representation) for each epitope group defined in Fig.2. CDR3 regions are colored in pink and FR3/FR4 anchors are colored in brown and light brown. (C) Dendrograms illustrating epitope similarity for IgGs and Nbs within each epitope group defined in Fig.2. Red lines at 0.4 (40% shared epitope residue identity) indicate the clustering threshold. Antibodies with a minimum pairwise epitope similarity of ≥ 0.4 are grouped into the same sub-cluster. (D) Table showing the number of sub-clusters and the percentage of antibodies contained in the three largest sub-clusters for each Nb and IgG epitope group. (E) Left: RBD surface contact propensity by IgG for the three largest epitope sub-clusters within IgG group IV (IgG sub-cluster IV.1, IV.2, and IV.3), comprising 11, 8, and 7 IgGs. Center: Superpositions of CDR3 loops (Cartoon representation) in complex with the RBD (surface representation). Right: Superpositions of CDR3 loops (Cartoon representation). The conformational outlier CDR3 loops (both IgGs and Nbs) are in white. (F) Left: RBD surface contact propensity by Nb for the three largest epitope sub-clusters within Nb group IV (Nb sub-cluster IV.1, IV.2, and IV.3), comprising 9, 8, and 6 Nbs. Center: Superpositions of CDR3 loops (Cartoon representation) in complex with the RBD (surface representation). Right: Superpositions of CDR3 loops (Cartoon representation). (G) Comparison of average pairwise CDR3-H sequence similarity within and between sub-clusters IgG VH IV.1, IV.2, and IV.3, scaled from 0 to 1. Sequence alignment of CDR3-H and FR3/FR4 anchors shown for IgG VH IV.1. Residues in the CDR3 region are colored according to their physicochemical properties, while FR3/FR4 anchor residues are shown in grey. (H) Comparison of average pairwise CDR3 sequence similarity within and between subclusters Nb IV.1, IV.2, and IV.3, scaled from 0 to 1. Sequence alignment of CDR3 and FR3/FR4 anchors shown for Nb IV.1. Residues in the CDR3 region are colored according to their physicochemical properties, while FR3/FR4 anchor residues are shown in grey.

We focused on the three largest sub-clusters within each epitope group, which collectively represent the majority of antibodies in each group, except Group II IgGs (25%) (**Fig. 3D**). Within these dominant sub-clusters, structural alignment of CDR3 loops revealed surprising conformational similarity (**Fig. 3E–F, fig. S6A–B**), despite minimal sequence identity (**Fig. 3G–H, fig. S6C–D**). These shared loop geometries were observed not only within sub-clusters targeting variable epitopes (**fig. S6**), but also across those targeting conserved regions (**Fig. 3E–F**). While convergence was evident within each antibody type, cross-species comparisons revealed substantial geometric divergence. Notably, Nbs consistently formed longer, more open CDR3 loops stabilized by a characteristic kinked conformation (*49*) (**Fig. 3F, fig.S6B**), likely an adaptation to expand paratope surface area without a VL. These observations illustrate a dual theme: antibody repertoires exhibit marked sequence and structural diversity, yet converge on shared paratope geometries tailored to specific epitopes. This convergence, emerging independently across distinct mammalian antibody species, indicates underlying geometric and physicochemical restraints that guide antigen recognition toward a limited set of structurally favorable solutions.

### Widespread and extensive epitope mutations

SARS-CoV-2 evolution has produced successive waves of variants capable of evading both vaccines and neutralizing antibodies (*50–52*). Over 30 variant-associated mutations have been identified in RBD. These mutations have not only extensively remodeled the RBS but also accumulated significant mutations in conserved regions, enhancing immune escape while optimizing receptor engagement (*53*) (**Fig. 4A and 4B**).

**Fig. 4:**
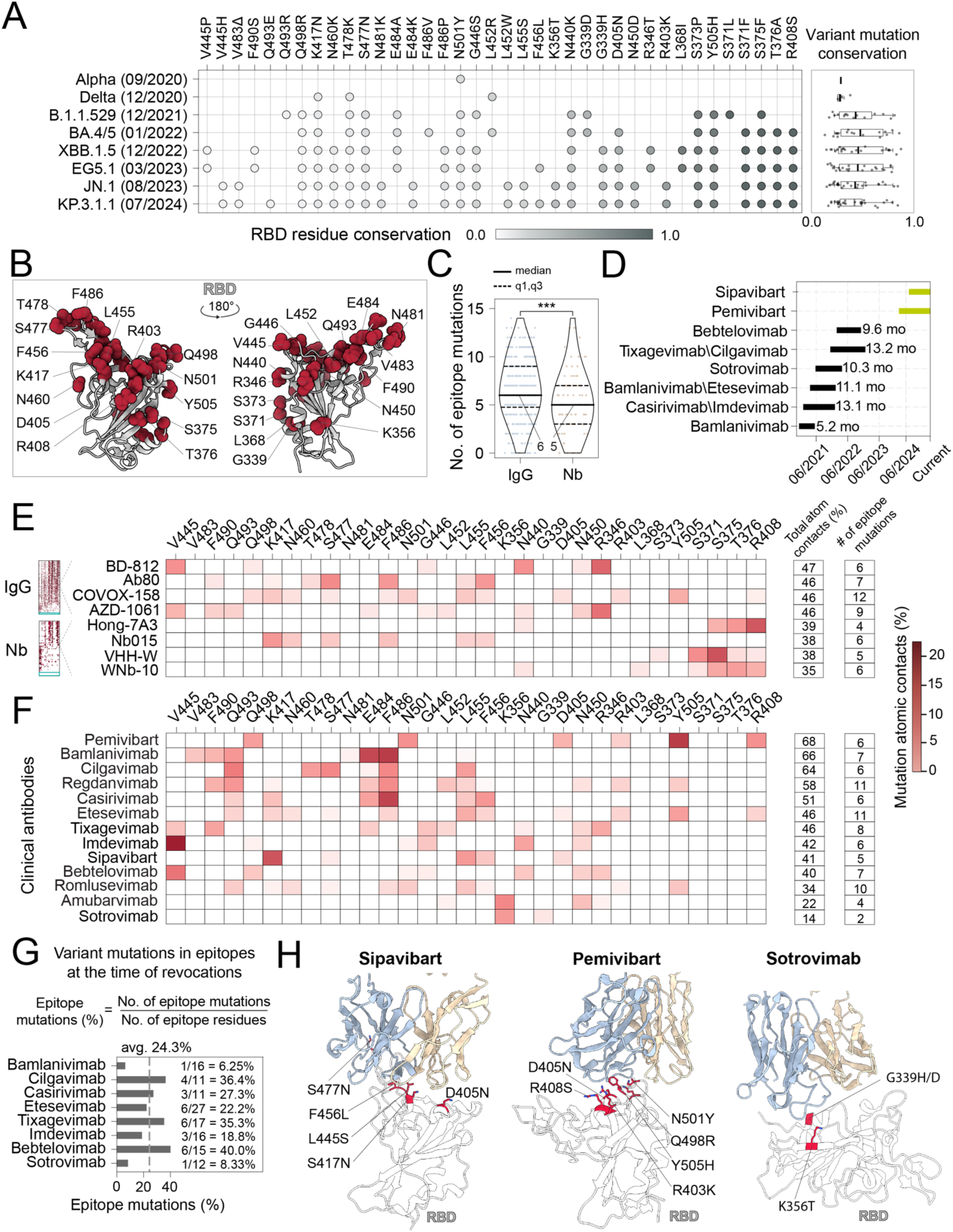
Epitope mutational landscape of RBD-targeting antibodies. (A) Mutations from different circulating variants on RBD. Left: A table shows the names of circulating variants in chronological order, mutations are sorted and colored based on corresponding wildtype residue conservation. Right: Boxplots showing conservation distributions of mutation sites for different circulating variants. The bold line represents the median, and the dashed lines represent the 25th, 75th percentile. (B) RBD (PDB:6M0J) is shown in cartoon representation, with mutations shown as red spheres. (C) Distribution of IgGs and Nbs with different numbers of mutation sites of circulating variants in their epitopes. (D) Timeline of emergency use authorization (EUA) and revocation for FDA and EMA approved antibodies. Sipavibart and Pemivibart are not officially revoked when this paper was drafted. (E) Left: Heatmaps showing percentage of atom contacts with mutation sites on epitopes for IgGs and Nbs, ranked by the total percentage of epitope mutation contacts from top to bottom. Cyan colored boxes indicate the proportion of antibodies without epitope mutation sites: 2.9% for IgG and 8.3% for Nb. Middle: Zoom-in view showing atom contacts for 4 representative IgGs and Nbs ranked in the middle. Right: tables showing the percentage of contacts to and number of mutation sites on epitopes for representative antibodies. (F) Heatmaps showing percentage of atom contacts with mutation sites for clinically approved IgG antibodies. (G) Percentage of epitope residues mutated in circulating SARS-CoV-2 variants at the time of clinical mAb revocation. Variant designations correspond to major lineages present during each FDA revocation decision: Bamlanivimab (Delta), Casirivimab, Imdevimab, and Etesevimab (BA.2.12.1), Sotrovimab (BA.5), and Tixagevimab, Cilgavimab, and Bebtelovimab (XBB.1.5). Mutation percentages reflect the extent of epitope remodeling associated with loss of neutralization potency. (H) Complex structures of the two most recently approved clinical mAbs, along with one targeting a moderately conserved region: Sipavibart (PDB: 8SUO), Pemivibart (PDB: 7U2D), and Sotrovimab (PDB: 7R6W). RBD mutation sites are colored in red

Our analysis reveals that more than 96% of antibody epitopes (97.1% for IgGs and 91.7% for Nbs) overlap with at least one mutation found in major circulating variants (**Fig. 4E, Table S1**). Only 3.9% of antibodies (2.9% of IgGs and 8.3% of Nbs) target entirely mutation-free epitopes (**Table S1, fig. S7B, S7D**). IgGs and Nbs interact with six and five mutated RBD residues (median, respectively (**Fig. 4C**), representing 40.3% and 30.6% of their epitope footprints (**fig. S7A**). All thirteen clinically authorized monoclonals show extensive overlap with mutated residues (**Fig. 4F–H, fig. S8, Table S1**). These mAbs typically recognize six mutated residues per epitope (median), corresponding to 24.3% of epitope sites, with the number ranging from 2 to 11 mutations (6.3%–40%) at the time of regulatory revocation (**Fig. 4G**).

Notably, 11/13 mAbs bind to the highly variable RBS (Groups I and II), while only Sotrovimab and Amubarvimab target moderately conserved regions (Group III; **fig. S9A–C**). Consequently, they showed short functional lifespans, from 5.2 months (Bamlanivimab) to 13.2 months (the Tixagevimab-Cilgavimab cocktail), before losing efficacy against emerging variants and ultimately being revoked (**Fig. 4D**). Moreover, the two most recently approved mAbs Pemivibart (ADG-2) and Sipavibart (authorized in March and July 2024, respectively) for prophylaxis in immunocompromised individuals, now exhibit either a >400-fold reduction in neutralization potency or complete loss of activity against the dominant Omicron KP.3 sublineages (KP.3.1.1 or KP.3.3) (*54, 55*). Similar escape has been observed for mAbs authorized in East Asia, although their formal withdrawal remains unconfirmed.

### Rare antibody survivors against highly immune-evasive Omicron variants

We curated a dataset of 72 IC_50_ measurements for RBD-targeting IgGs and Nbs across wild-type SARS-CoV-2 and major variants (*56, 57*) (**Table S3**), to evaluate whether structural features of antibody–antigen interfaces could predict loss of neutralization against emerging variants. For each antibody, we quantified epitope disruption using two metrics: mutated epitope residues and the number of atomic contacts disrupted by variant-associated mutations (**Methods**). We found a strong positive correlation between the log_10_-fold change in IC_50_ and the percentage of mutated epitope residues (Pearson r = 0.68; **fig. S10C**), indicating that antibodies targeting more frequently mutated surfaces tend to lose potency. This correlation is considered particularly strong considering significant variability across experimental conditions and assay platforms (**fig. S10B, Table S4**), and was slightly stronger for Nbs (r = 0.71) than for IgGs (r = 0.66). Molecular dynamics simulations further supported these observations: for Nbs, predicted reductions in contact surface area correlated strongly with diminished potency (R = –0.78), whereas this trend was less consistent for IgGs (**fig. S10A, S10D, Table S5**).

Nbs preferentially target conserved, high-fitness epitopes, implying they may be more resilient against mutations. Their robust solubility and stability also facilitate reproducible experimental assessments. From an initial set of 20 PDB-derived Nbs with fewer than 15% overall epitope mutations (**Fig. 5A, Table S1**), we prioritized the top 12 candidates that have not shown substantial loss of neutralization against tested variants (*42, 44, 57–59*). These selected Nbs were robustly expressed in E. coli and evaluated by ELISA for binding to recombinant RBDs from the WT virus, Omicron BA.2, and KP.3 (*60*). Their neutralizing activity was measured using an authentic virus microneutralization assay (*61, 62*).

**Fig. 5:**
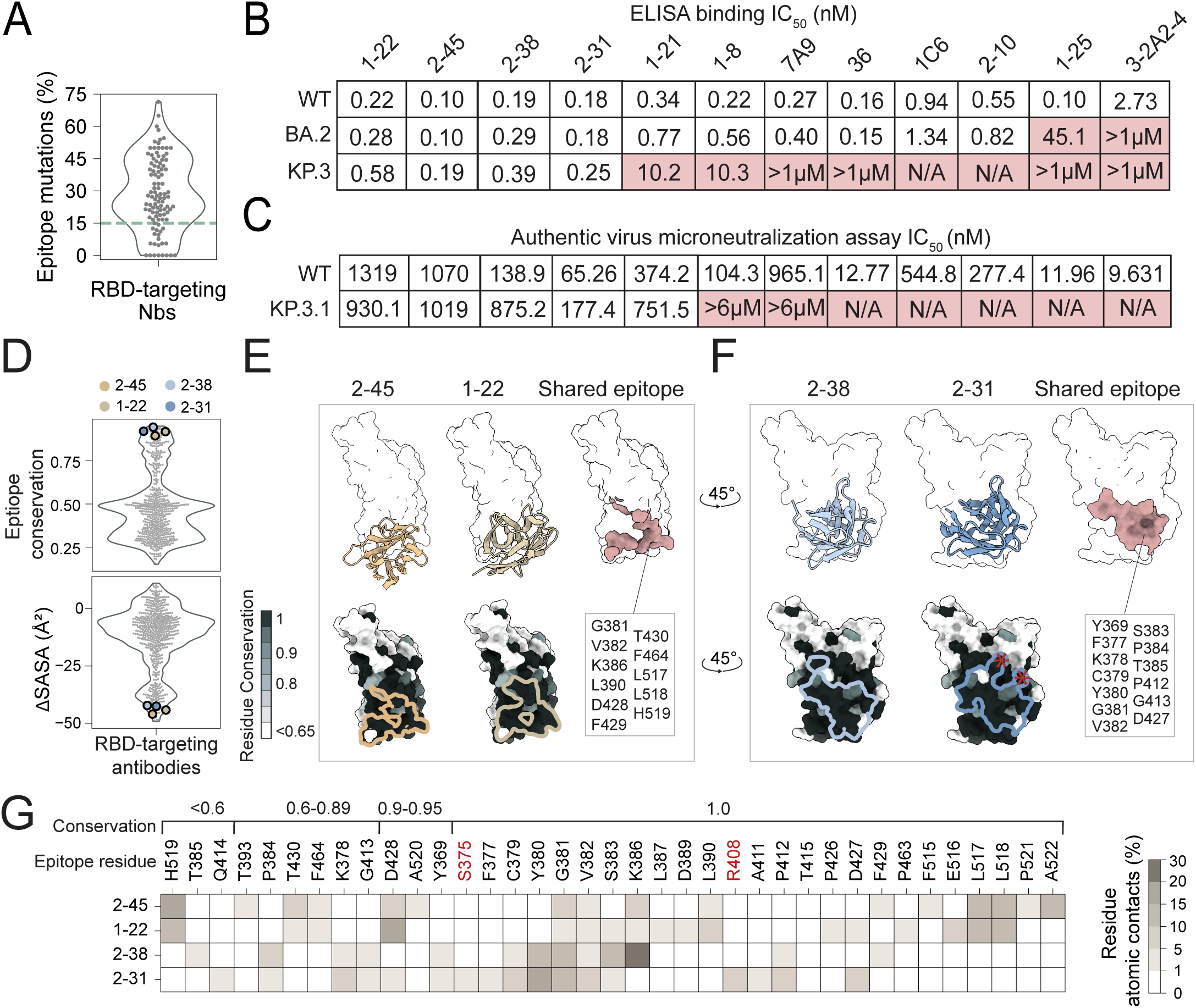
Evaluation of top 12 RBD Nbs from PDB against the advanced Omicrons. (A) Violinplot showing distribution of percentage of epitope mutations for Nbs. The green dashed line indicates the 15% epitope mutation cutoff used for Nb selection. (B) Summary of binding affinity to WT, BA.2 and KP.3 RBDs by ELISA of selected Nbs. N/A: No binding at the highest tested concentration (10 µM). Binding affinity highlighted in read indicates > 10-fold decrease by the variants compared to the WT. Note KP.3 and KP.3.1.1 share the identical RBD sequence. (C) Neutralization potency of selected Nbs against the authentic SARS-CoV-2 viruses. Colored in red: More than 10-fold decrease in neutralization compared to the WT. KP.3.1 and KP.3.1.1 share an identical RBD sequence. (D) Violin plots showing distribution of epitope conservation(upper) and epitope accessibility on Spike (Δ SASA) for RBD-targeting antibodies. Nbs that retain strong binding and neutralization of KP.3.1 are colored in circles. (E-F) Epitope characteristics of four broad-spectrum Nbs. Upper: Nbs (cartoon representation) in complex with RBD (surface representation), shared epitope is colored in salmon and exact residues are shown. Lower: RBD is colored by residue conservation, the corresponding epitope is outlined. Epitope mutations are labeled in star. (G) Heatmap showing the percentage of atom contacts of Nbs and their epitopes. The epitopes residues are sorted by conservation. Mutations are in red.

10/12 Nbs retained strong binding to BA.2, and only four (1–22, 2–45, 2–38, 2–31) maintained robust binding to the KP.3 RBD (**Fig. 5B, fig.S11**). These four converged on two epitopes (Groups III and IV) that are among the most conserved and cryptic identified to date, bearing minimal if any mutations (**Fig. 5D-G**). Nbs 1-22 and 2-45, despite strong binding, exhibited weak neutralization (∼1 µM IC_50_) against both WT and KP.3.1 (**Fig. 5C, fig.S12**). In contrast, Nbs 2-38 and 2-31 achieved moderate neutralization of KP.3.1, with IC_50_s of 875 nM and 177 nM, respectively, reflecting <10-fold reductions from WT potency. Collectively, these findings underscore the outstanding challenges in identifying antibodies that can effectively neutralize highly evolved variants. Long-lived, mutation-resilient antibodies like Nb 2-31 are likely very rare with compromised potency.

## Discussion

The rapid evolution of SARS-CoV-2 has profoundly shaped global health and biomedical research, prompting an unprecedented international effort to develop effective countermeasures—including vaccines, small-molecule inhibitors, and therapeutic antibodies(*2*). Upon infection or vaccination, the adaptive immune system generates a diverse repertoire of antibodies capable of recognizing viral proteins with remarkable specificity and affinity. These antibodies neutralize virions directly and engage host effector functions to promote viral clearance(*9, 12, 15, 63*).

Understanding the molecular basis of antibody recognition against evolving viral antigens lies at the heart of immunology and structural biology(*31*). To date, over a thousand of antibody–antigen structures have been resolved, representing a milestone in structural immunology and creating a powerful resource for dissecting the principles that govern antibody targeting and immune escape. While most previous studies focused on individual antibodies, our analysis enables a systems-level view of how the mammalian immune repertoire recognizes, adapts to, and is constrained by viral evolution.

Several key insights emerge from our analysis. First, we find that virtually every surface residue of the viral RBD is antigenic (**Fig. 2E**), despite consistent binding preferences for specific epitope regions (**Fig. 2F**). This widespread antigenicity underscores the versatility of mammalian antibodies and suggests broad immune pressure across the RBD surface, potentially contributing to rapid and continuous antigenic drift. These patterns mirror those seen in other rapidly evolving pathogens, such as influenza and HIV (*64–66*).

Despite significant variation in paratope composition, we observe striking structural convergence across antibodies targeting the same epitope clusters (**Fig. 3E, 3F, fig. S6A, S6B**). This convergence, observed in both IgGs and Nbs, suggests that antibody recognition is funneled through a limited set of biophysically favorable binding modes(*67, 68*). These modes likely reflect shared constraints in shape complementarity and electrostatics that govern productive antigen engagement. Thus, a dual principle emerges: remarkable sequence and structural diversity, yet convergence on optimal paratope configurations tailored to specific epitopes.

Comparative analysis further reveals distinct epitope-targeting strategies. IgGs predominantly recognize large, solvent-accessible regions, particularly within the RBS that are prone to variation. In contrast, Nbs tend to engage smaller, conserved, and often cryptic epitopes using compact paratopes and enhanced electrostatic interactions(*69*). These differences imply structural adaptations that account for the greater mutational tolerance observed in select Nbs. Critically, our analysis shows that antibodies, including all clinically approved monoclonals, target epitopes that overlap with multiple variant-associated mutations, affecting over one-third of their contact residues. As a result, most show sharply reduced or abolished activity against highly immune evasive variants such as Omicron KP.3.1.1(*54, 60, 70*). This is consistent with >170 RBD antibodies that we have been able to find from literature: virtually all have been evaded by waves of variants and no potent neutralizer of KP.3.1.1 has yet been documented (**Table S6, fig. S14**). Even Nbs that bind the most conserved epitopes fail to neutralize potentially due to structural inaccessibility (**fig. S13**) and/or a lack of receptor competition. These findings highlight intrinsic trade-offs in single-epitope strategies, underscoring the outstanding challenges in sustaining protection amid rapid antigenic evolution (**fig. S14**).

In sum, our findings reveal general principles of antibody recognition, structural convergence, and immune escape that extend beyond SARS-CoV-2. By integrating large-scale structural mapping, computational epitope profiling, and experimental validation, we establish a framework for understanding how mammalian immune systems adapt to viral evolution under constraint. This approach informs the rational design of next-generation antibodies that harness multi-epitope engagement and engineered avidity(*57, 71*), which are essential for countering immune escape and enhancing pandemic preparedness. While this study focuses primarily on RBD-directed antibodies, which reflects their immunodominance and structural tractability, our analysis reveals similar escape patterns across other spike regions such as the NTD(*13*) (**fig. S7C, S7E, S7F**). Antibodies targeting more conserved S2 regions, including the fusion peptide and stem helix, remain rare and typically show limited potency, despite their promise as broadly protective targets(*9, 13, 72–75*). Expanding structural and functional characterization across underexplored epitopes will help build a more complete view of antiviral humoral immunity.

## Supplemental Information

**Supplementary Fig. 1.**
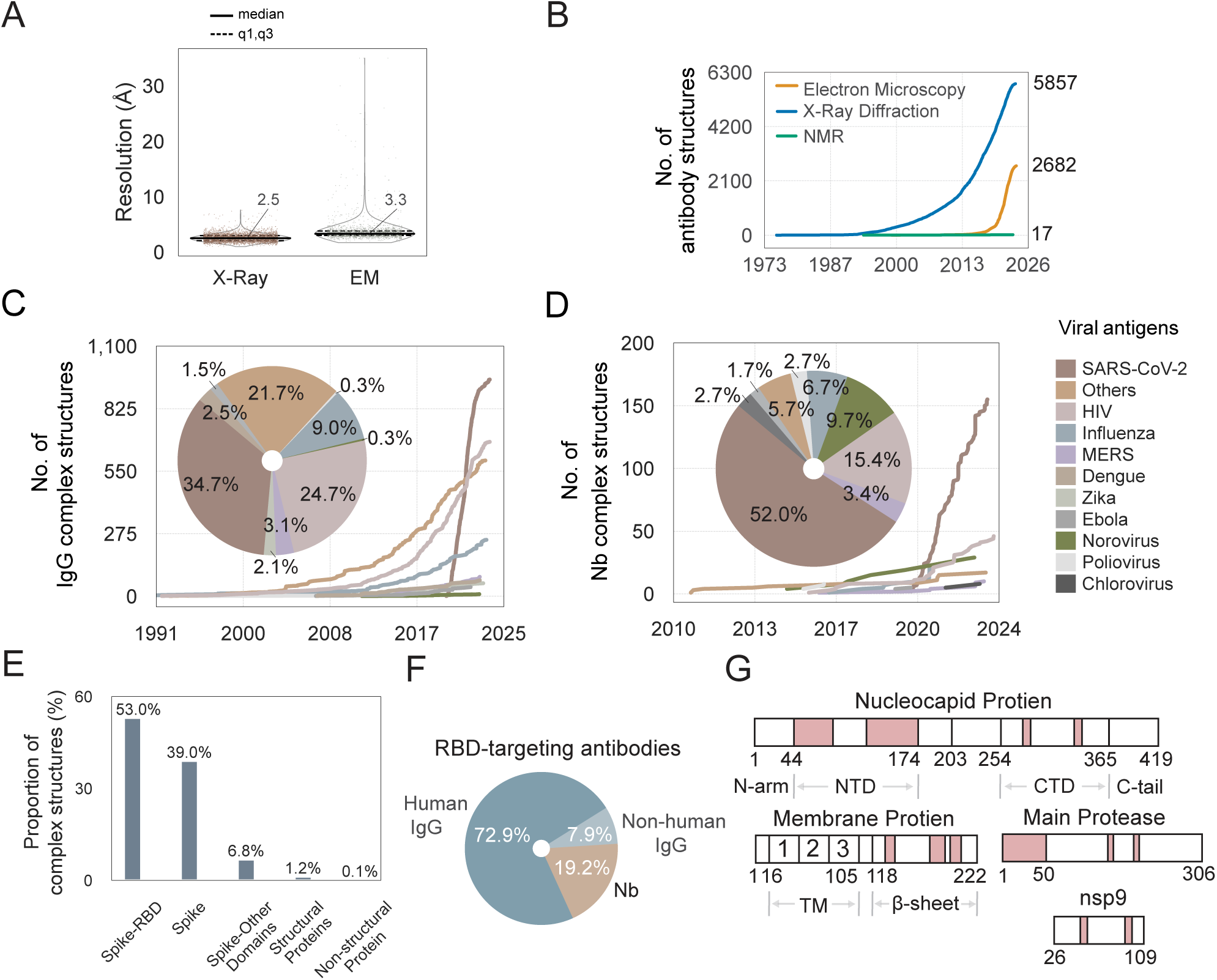
Antibody complexes targeting SARS-CoV-2 functional proteins: (A) Violin plots showing distribution of resolution of antibody-antigen complex structures determined by X-ray crystallography and cryoEM. (B) Cumulative count of antibody structures categorized by structure determination methods. (C-D) Cumulative count of IgG/Nb structures in complexes with different viral proteins. The inset pie chart shows the percentage of individual viruses. (E) Distribution of antibody structures (1,108 unique PDB entries) in complex with different SARS-CoV-2 antigens. (F) Pie chart showing percentage of IgG and Nb targeting SARS-CoV-2 RBD. (G) Schematic representation of the SARS-CoV-2 functional proteins targeted by antibodies, including nucleocapsid protein, non-structural protein 9 (nsp9), membrane protein and main protease. Antibody epitopes are colored in pink.

**Supplementary Fig. 2.**
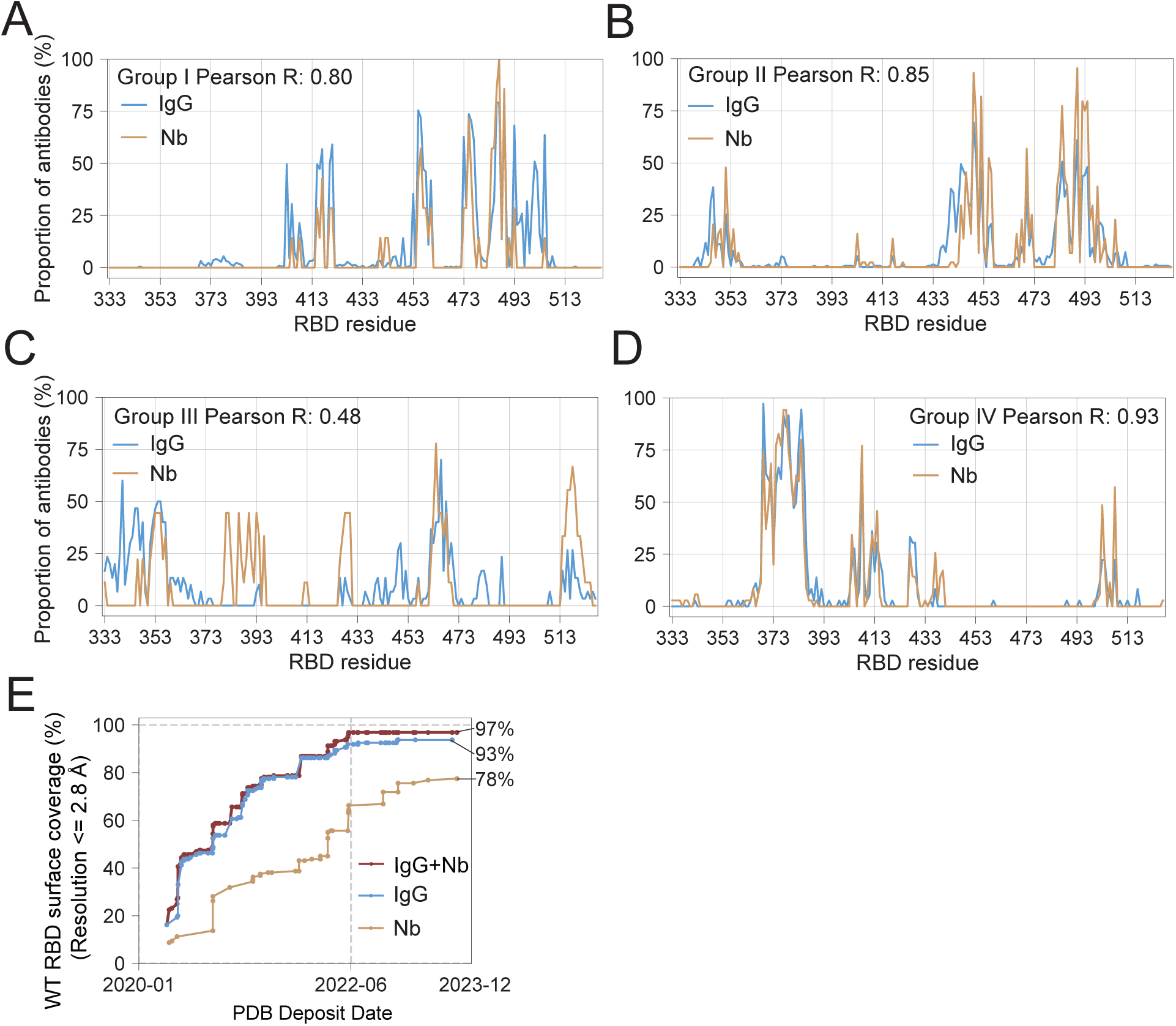
Epitope classification and coverage: (A-D) Comparison of residue binding propensity between IgGs and Nbs in each epitope group, Pearson correlation is calculated. (E) Cumulative WT RBD surface coverage by antibodies from structures with resolution < 2.8 Å.

**Supplementary Fig. 3.**
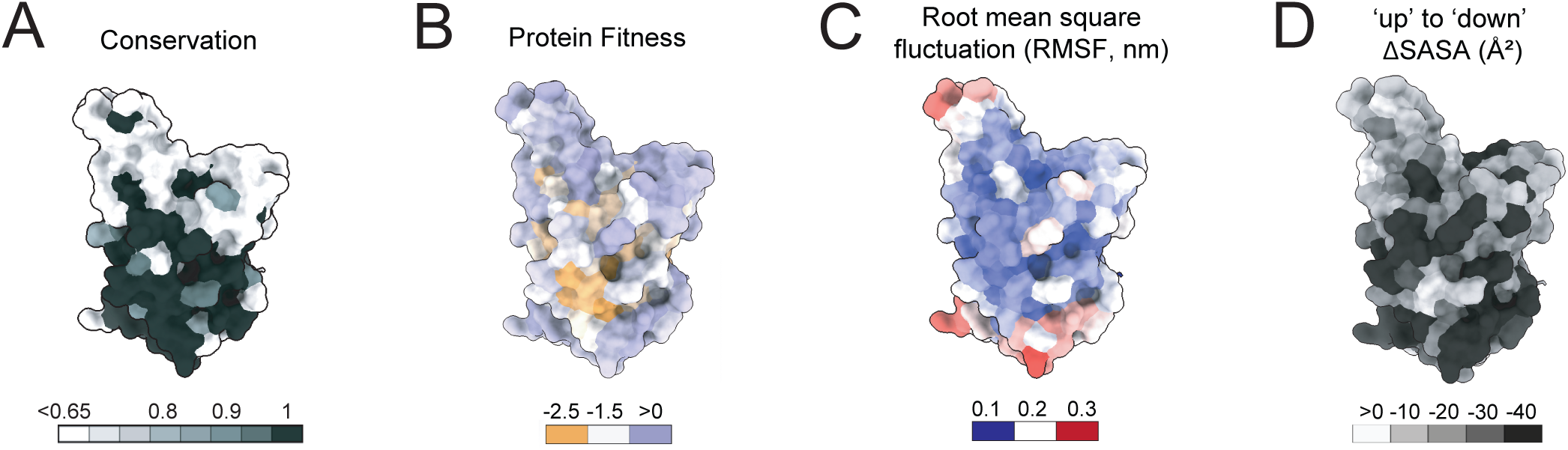
Epitope properties of RBD antibodies: (A) RBD surface representation showing sequence conservation based on 19 sarbecoviruses. (B) RBD surface representation showing protein fitness based on mutational effects on RBD expression level. (C) RBD surface representation colored by structural flexibility based on root-mean-square-fluctuation (RMSF) from MD simulation. (D) RBD surface representation showing residue ΔSASA from RBD in ‘up’ conformation to ‘down’ conformation.

**Supplementary Fig. 4:**
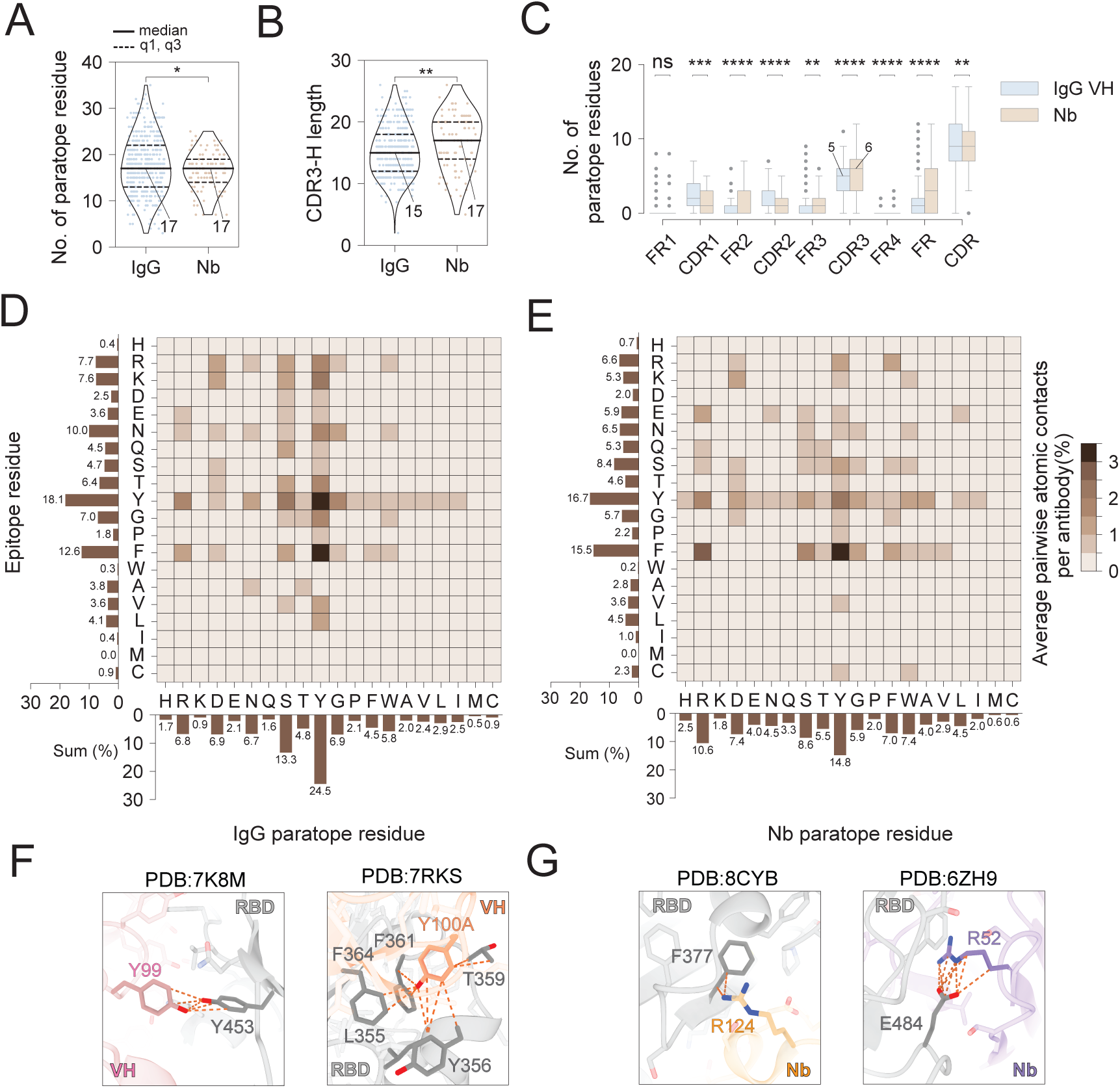
RBD interactions by IgGs and Nbs. (A) Comparison of number of paratope residues between IgGs and Nbs. The bold line represents the median, and the dashed lines represent the 25th, 75th percentile. Statistical analysis was performed using a two-tailed student t test, \**p* < 0.05, \*\**p* < 0.01, \*\*\**p* < 0.001, \*\*\*\**p* < 0.0001. (B) Comparison of number of residues in heavy chain CDR3 between IgGs and Nbs. (C) Box plots comparing the number of paratope residues between IgGs and Nbs from different FRs and CDRs. (D-E) Heatmaps showing pairwise interaction between epitope and paratope residues for IgGs (left) and Nbs (right). Bar plots showing the total average percentage of pairwise atomic contacts per antibody for each specific amino acid on the paratope or epitope. (F) Representative tyrosine-involved pi-pi interaction between IgGs and RBD. (F) Representative arginine-involved cation-pi (left) and electrostatic (right) interactions between Nbs and RBD.

**Supplementary Fig. 5.**
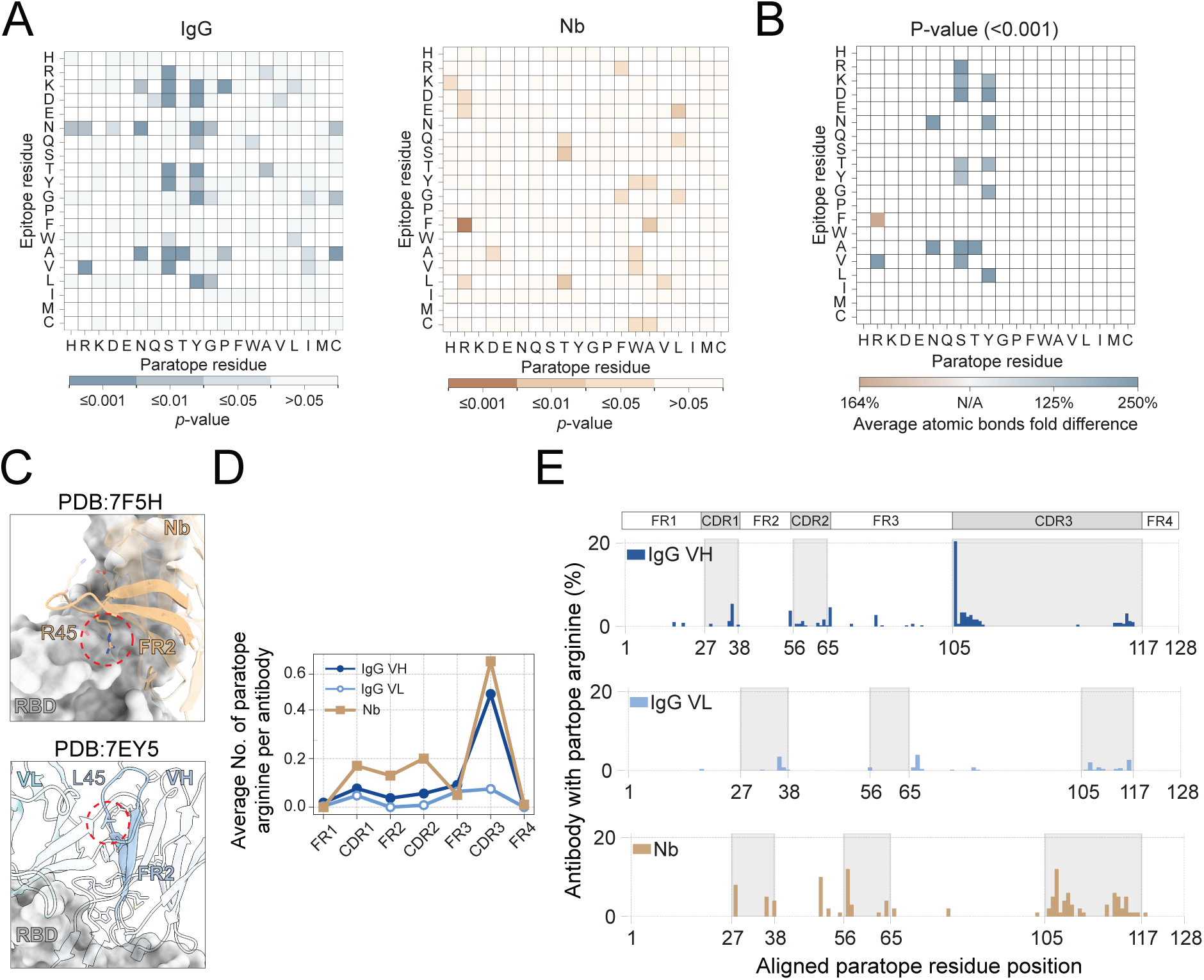
Analysis of molecular interactions between RBD and antibodies: (A) Heatmaps showing *p*-values comparing the average pairwise epitope-paratope atomic contacts per antibody between IgG and Nb. Left: *p*-values for prevalent pairwise contacts are shown for IgGs. Right: *p*-values for prevalent pairwise contacts are shown for Nbs. (B) Heatmap showing fold changes in average atomic contacts between IgG and Nb, only statistically significant (*p*-value < 0.001). (C) Upper: Representative Nb-RBD complex structure showing FR2-involved binding (R50 in IMGT scheme). Lower: Representative IgG-RBD complex structure showing heavy chain FR2 is buried by light chain (L50 in IMGT scheme). (D) Average number of paratope Arginine for each IgG and Nb chain by different FRs or CDRs. (E) Percentage of antibodies with arginine in the paratope at each aligned residue position (IMGT scheme). CDR regions are highlighted in grey.

## Summary of the results shown in Supplementary Fig.4 and 5

To dissect the molecular determinants of high-affinity binding, we analyzed paratope residues including contributions from complementarity-determining regions (CDRs) and framework regions (FRs). Nb paratopes comprise a median of 17 residues, comparable to 17 residues observed in IgG paratopes across both heavy and light chains. Nbs possess a significantly longer CDR3 loop which contributes more residues for interaction compared to IgG heavy chains (**Supplementary Fig. 4B,4C**), which may compensate for the absence of a light chain by expanding the binding interface(*69, 76*).In addition, we found that Nbs can uniquely leverage FR2 (**Supplementary Fig. 4C**), including a hallmark residue (e.g., R50 in IMGT numbering)(*76*), where the counterpart residue L50 is generally buried in IgG (**Supplementary Fig. 5C**), to contribute directly to antigen binding.

We next analyzed pairwise paratope–epitope interactions and generated residue contact heatmaps (**Supplementary Fig. 4D and 4E, Methods**). In IgGs, tyrosine (Y) and serine (S) residues are markedly enriched in paratopes, each contributing 24.5% and 13.3%, respectively, of all observed interaction contacts (**Supplementary Fig. 4D**). Here paratope Y is often observed to insert into RBD pocket, forming a pi-pi interaction with other bulky aromatic epitope residues such as Y or phenylalanine (F) (**Supplementary Fig. 4F**). Although Y and S are also enriched in Nbs, their contributions are substantially lower, accounting for 14.8% and 8.6% of the overall contacts, respectively (**Supplementary Fig. 4E**). Instead, Nbs preferentially utilize electrostatic interactions for binding. Their paratope arginine (R) alone contributes 10.6% of total interaction contact, representing the most significantly overrepresented residue compared to IgGs (**Supplementary Fig. 4E, Fig. 5A and 5B**). Specifically, they frequently form cation-pi interactions between paratope arginine (R) and epitope phenylalanine (F), as well as salt bridges between R and glutamic acid (E) (**Supplementary Fig. 4G**), occur roughly at twice the frequency seen in IgGs (**Supplementary Fig. 4D and 4E**). We further located the paratope R and found that it spreads across all three CDRs and FR2 of Nbs (**Supplementary Fig. 5D**), specifically at positions 28, 50, 57, 108, and 111 based on the IMGT scheme (**Methods**). The R in IgG is specifically enriched at position 106 within the CDR3 (**Supplementary Fig. 5E**). Analysis of RBD binding across a large and diverse antibody panel aligns with prior studies of global antibody–antigen complexes with more limited sampling (*77*). Our findings support the view that antibody repertoires are evolutionarily tuned for high-affinity binding.

---

**Supplementary Fig. 6.**
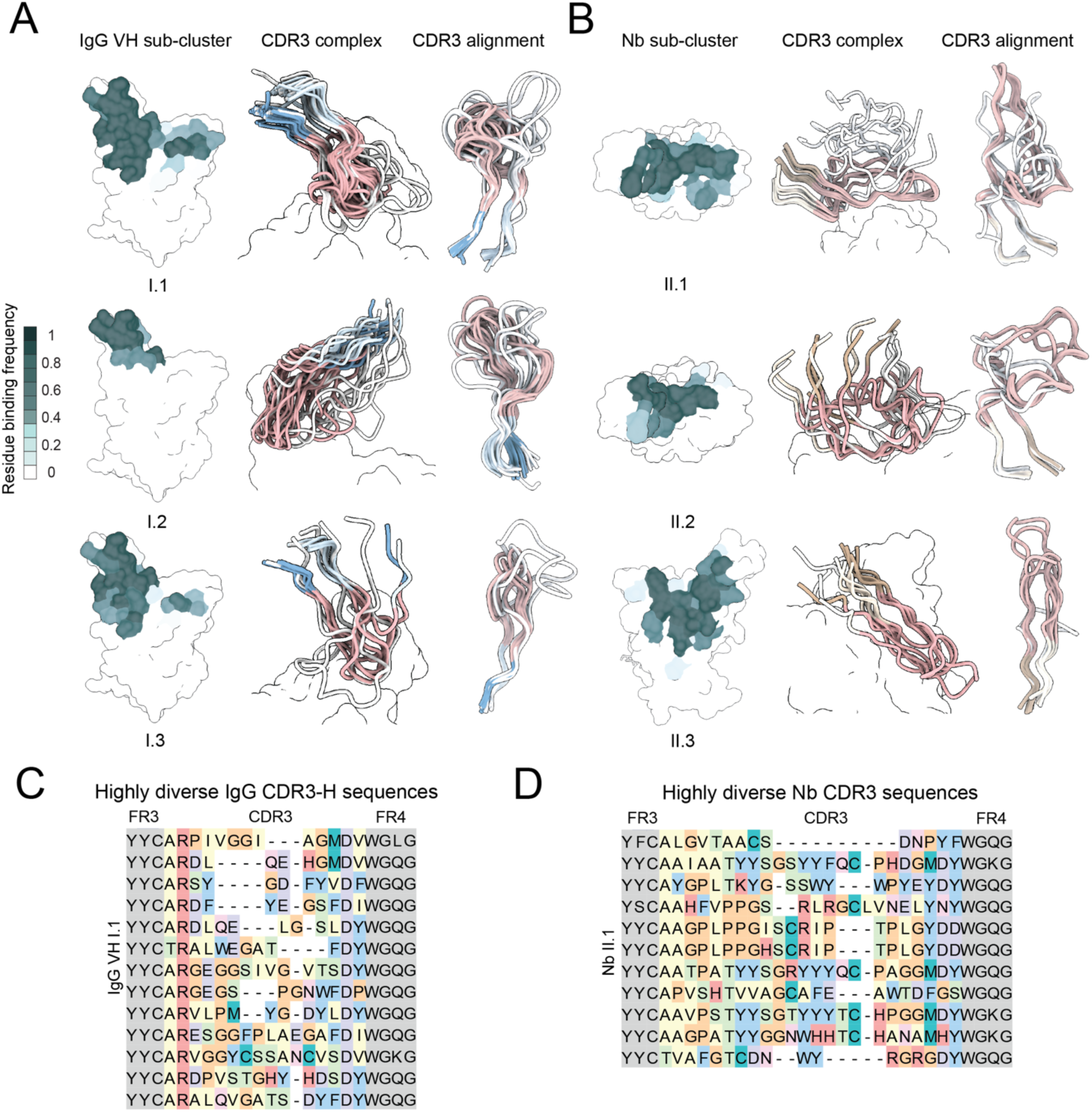
Epitope sub-clusters and CDR3 loop conformation convergence on the SARS-CoV-2 RBS: **(A)** Left: RBD surface contact propensity by IgG for the three largest epitope subclusters within IgG group I (IgG sub-cluster I.1, I.2, and I.3), comprising 81, 21, and 10 IgGs. Center: Superpositions of CDR3 loops (Cartoon representation) in complex with the RBD (surface representation). Right: Superpositions of CDR3 loops (Cartoon representation). CDR3 regions are colored in pink and FR3/FR4 anchors are colored in blue and light blue. **(B)** Left: RBD surface contact propensity by Nb for the three largest epitope subclusters within Nb group II (Nb sub-cluster II.1, II.2, and II.3), comprising 11, 8, and 7 Nbs. Center: Superpositions of CDR3 loops (Cartoon representation) in complex with the RBD (surface representation). Right: Superpositions of CDR3 loops (Cartoon representation). CDR3 regions are colored in pink and FR3/FR4 anchors are colored in brown and light brown. **(C)** Comparison of average pairwise CDR3-H sequence similarity within and between subclusters IgG VH I.1, I.2, and I.3, scaled from 0 to 1. Sequence alignment of CDR3-H shown for a subset of IgG VH I.1. Residues in the CDR3 region are colored according to their physicochemical properties, while FR3/FR4 anchor residues are shown in grey. **(D)** Comparison of average pairwise CDR3 sequence similarity within and between subclusters Nb II.1, II.2, and II.3, scaled from 0 to 1. Sequence alignment of CDR3 shown for Nb II.1. Residues in the CDR3 region are colored according to their physicochemical properties, while FR3/FR4 anchor residues are shown in grey.

**Supplementary Fig. 7.**
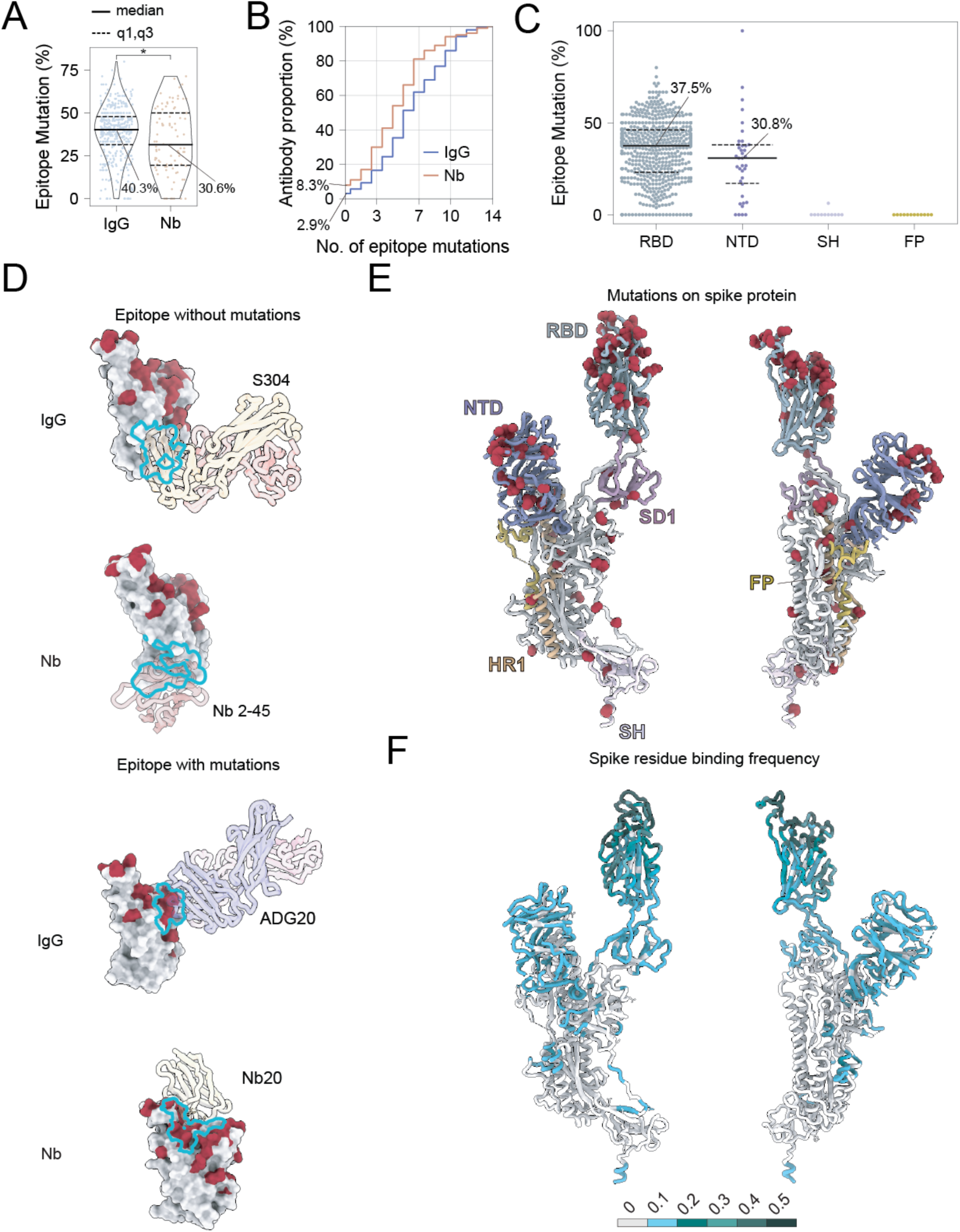
Epitope mutation and antibody binding on the SARS-CoV-2 spike protein: (A) Percentage of epitope mutations of circulating variants. The bold line represents the median, and the dashed lines represent the 25th, 75th percentile. Statistical analysis was performed using a two-tailed student t test, \**p* < 0.05, \*\**p* < 0.01, \*\*\**p* < 0.001, \*\*\*\**p* < 0.0001. (B) Cumulative percentage of IgGs and Nbs with increasing numbers of mutation sites of circulating variants in their epitopes. (C) Comparison of the percentage of epitope mutations of circulating variants in spike domains targeted by at least 10 antibodies, including the receptor binding domain (RBD), N-terminal domain, fusion peptide, and stem helix. (D) Schematics of epitope with mutations and without mutations for IgGs and Nbs. Representative structures include S304 (PDB: 8HWT), Nb 1-23 (PDB: 8CY9), ADG20 (PDB: 7U2D), and Nb 2-45 (PDB: 8CYD). (E) Structure of SARS-CoV-2 spike protomer is shown in cartoon representation, mutations from circulating variants are shown as red spheres, domains targeted by at least one antibody are colored. (F) Binding propensity of antibodies on spike residues (PDB: 6X2A).

**Supplementary Fig. 8.**
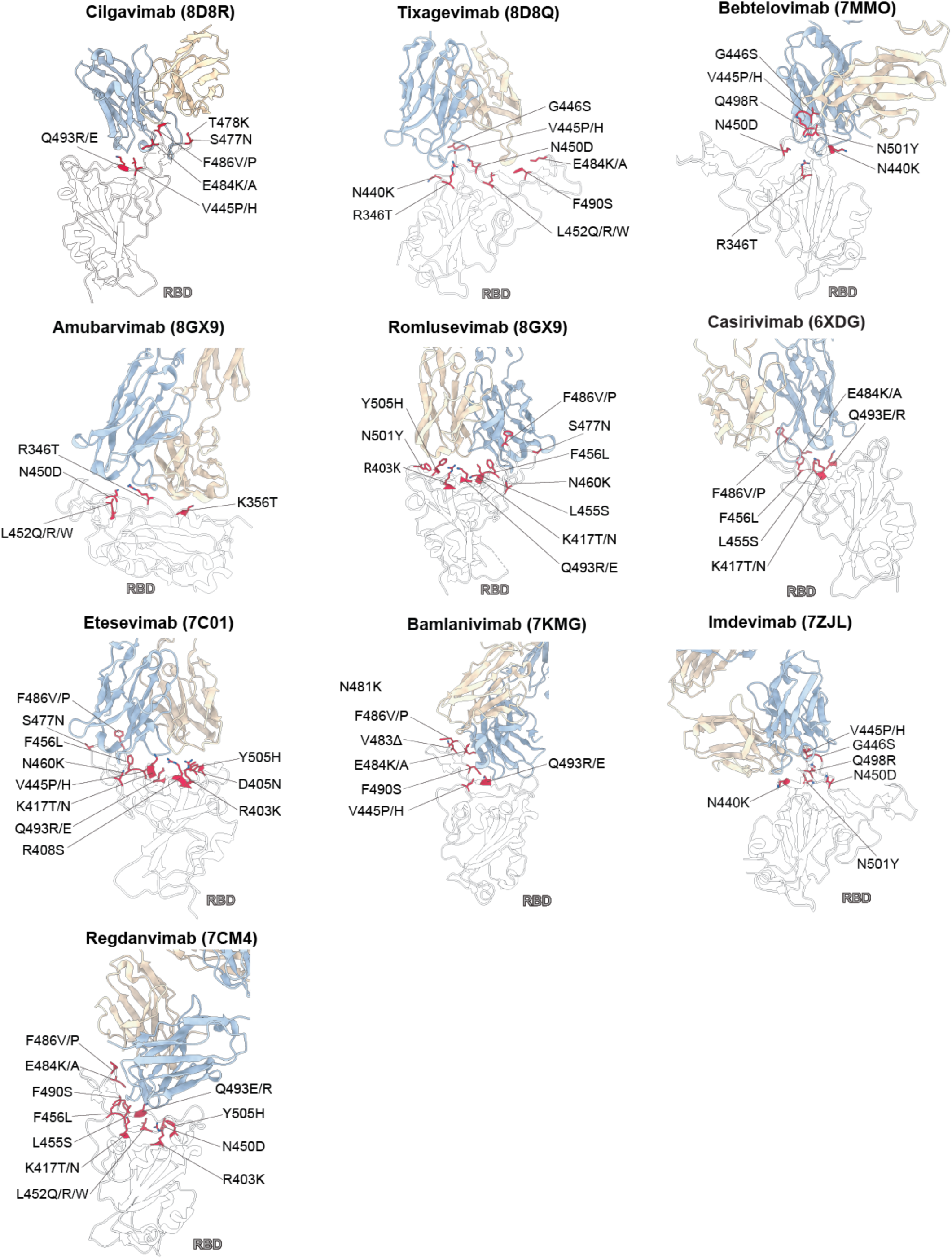
Structures of 13 clinical antibodies in complex with RBD. Residues mutated in the circulating variants are highlighted in red with side chain presented.

**Supplementary Fig. 9.**
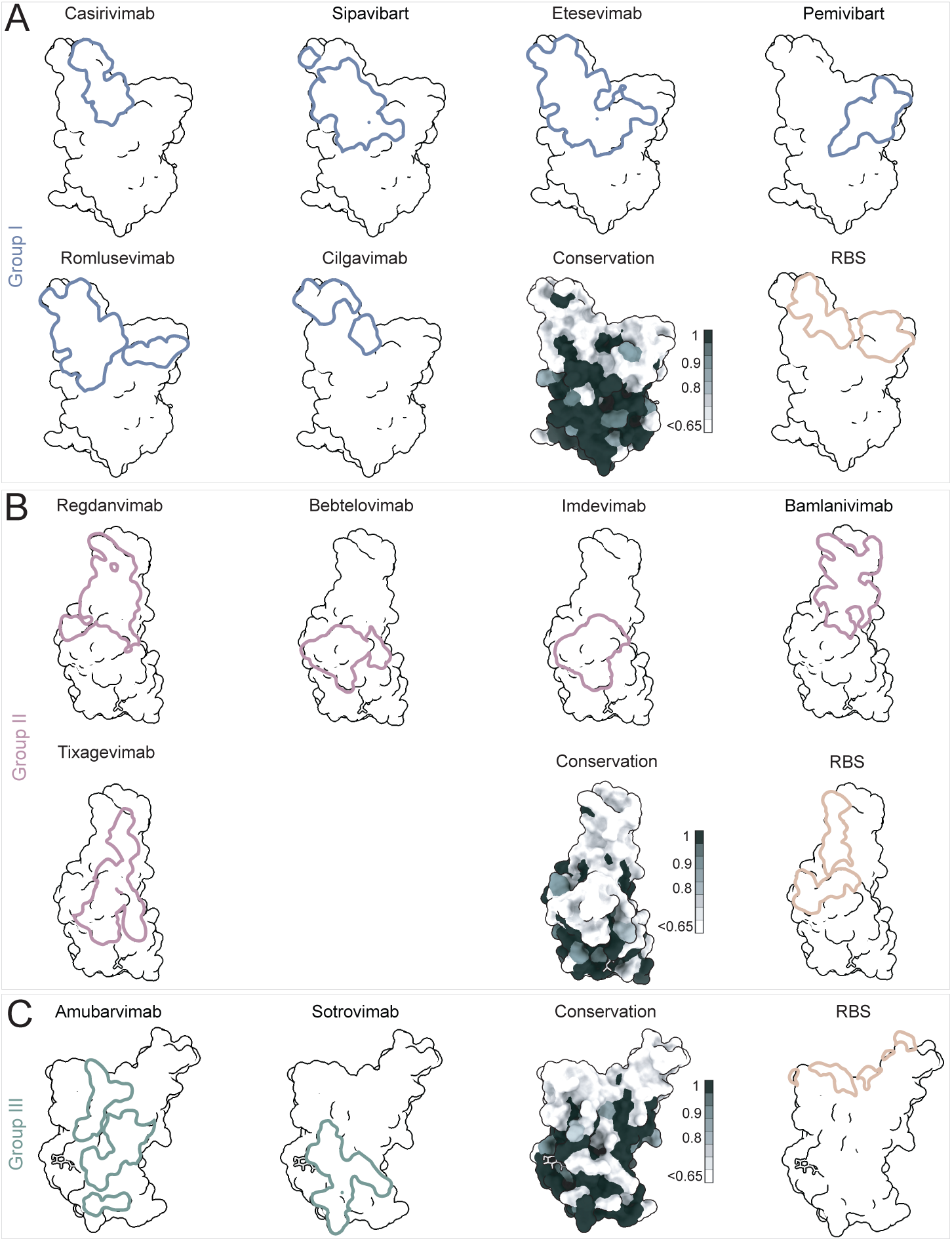
Epitopes of clinical antibodies: (A) RBD surface representation highlighting the epitope of clinical antibodies classified as Group I, shown alongside additional RBD surfaces colored by sequence conservation and RBS residues. (B) RBD surface representation highlighting the epitope of clinical antibodies classified as Group II, shown alongside additional RBD surfaces colored by sequence conservation and RBS residues. (C) RBD surface representation highlighting the epitope of clinical antibodies classified as Group III, shown alongside additional RBD surfaces colored by sequence conservation and RBS residues.

**Supplementary Fig. 10.**
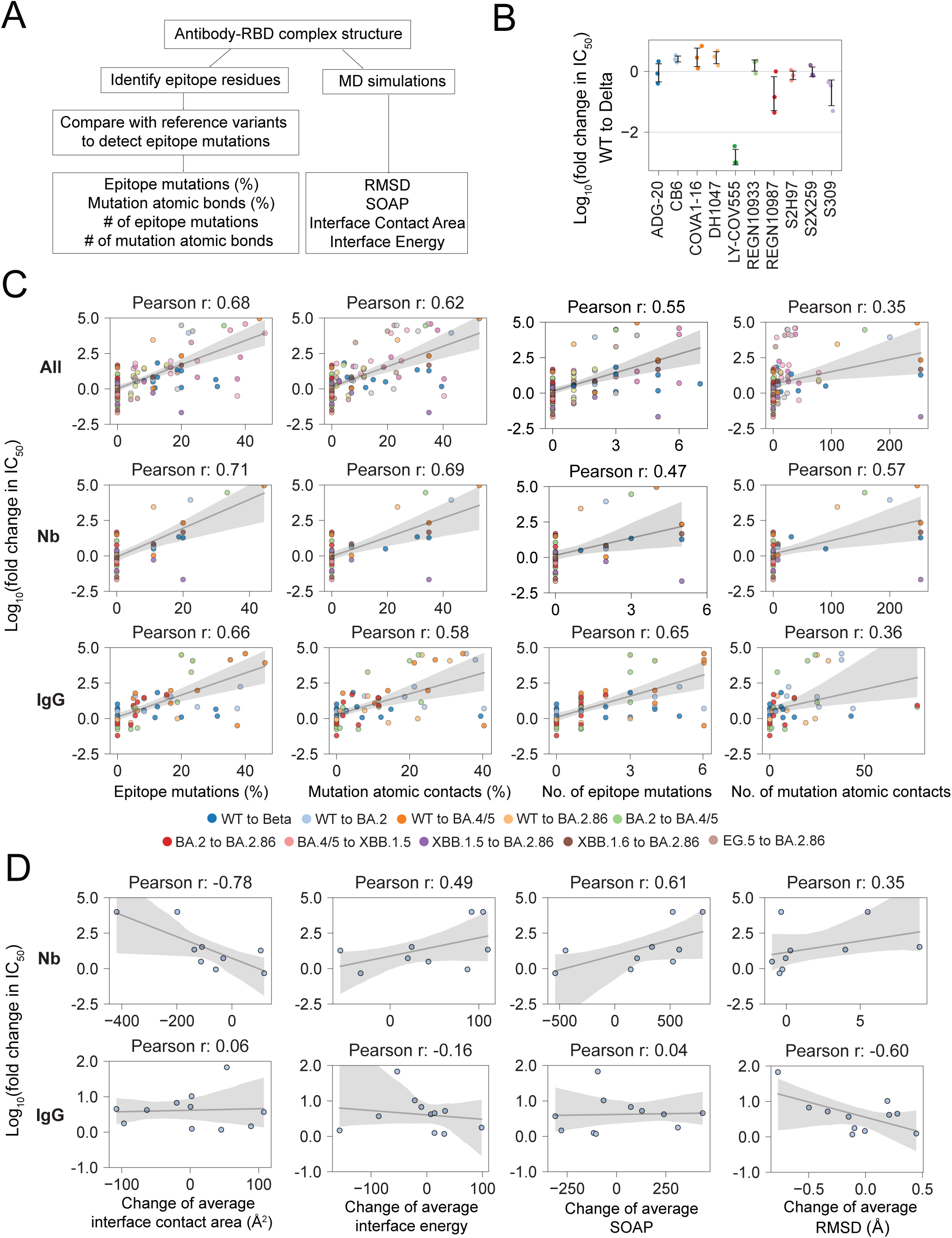
Correlating antibody neutralization efficacy across SARS-CoV-2 variants with mutational and simulated structural metrics: (A) Schematics of the workflow for correlating neutralization efficacy of antibodies against SARS-CoV-2 variants with by structure analysis and MD simulation. (B) Distribution of log_10_ fold change in IC_50_ from SARS-CoV-2 WT to Delta variant published by 3 different labs for each antibody. Error bar showing mean ± one standard deviation. (C) Correlation between log_10_ fold change in IC_50_ across SARS-CoV-2 variants and changes of metrics based on the RBD variant in their complex structures (percentage of epitope mutations, percentage of atomic contacts at mutated sites, number of epitope mutations and number of atomic contacts at mutated sites). Upper: correlation for the full dataset (IgGs and Nbs), center: correlation for the Nb dataset, lower: correlation for the IgG dataset. Each color represents a measure between two variants. Linear regression line shown with a 95% confidence interval. (D) Correlation between log_10_ fold change in IC_50_ from WT to BA.2 variant and changes of average MD simulation metrics (interface contact area, interface energy, SOAP score, and RMSD). Upper: correlation for the Nb dataset, lower: correlation for the IgG dataset.

**Supplementary Fig. 11.**
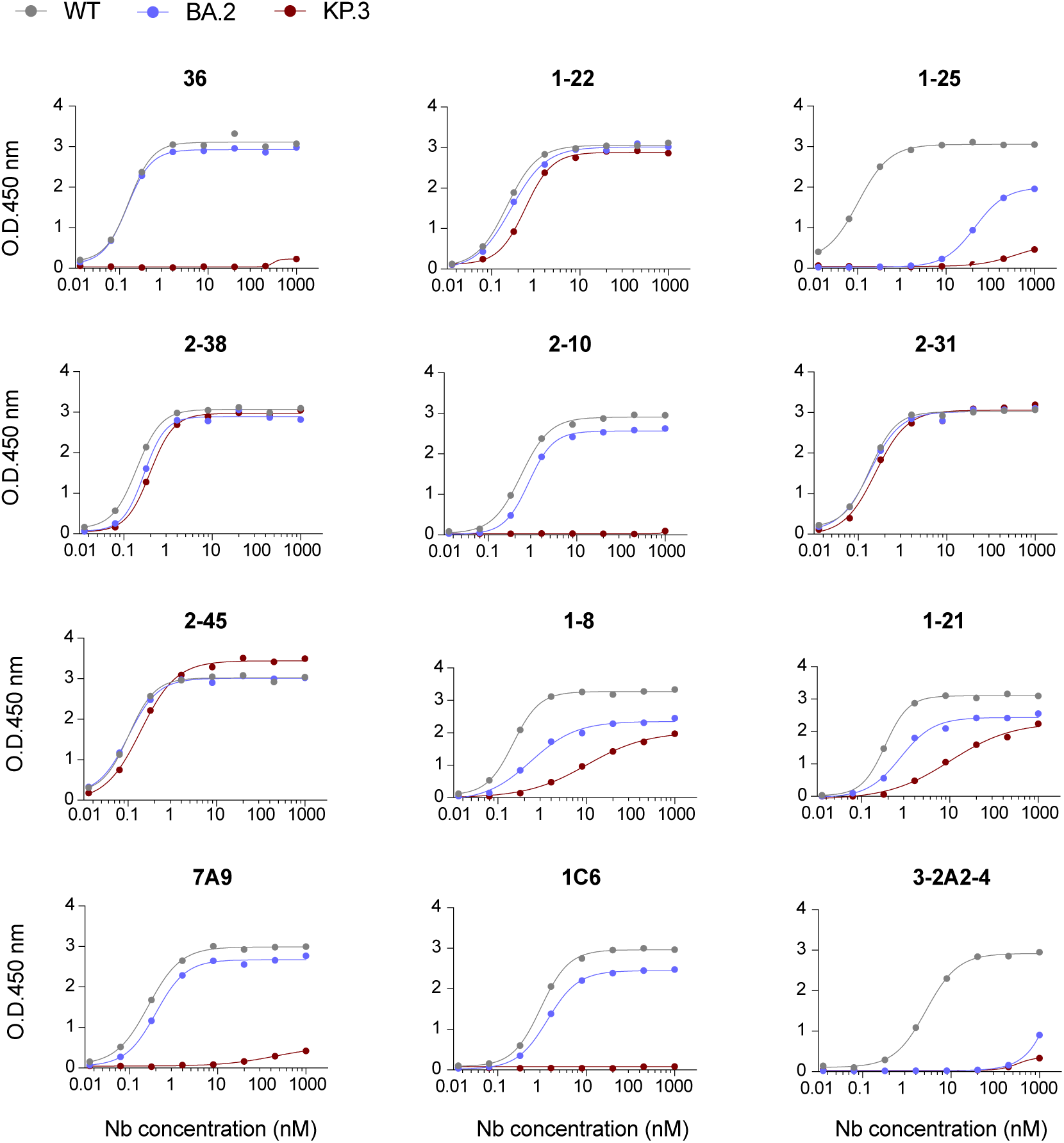
ELISA analysis of RBD Nbs: ELISA binding of top 12 RBD Nbs (associated with < 15% of epitope mutated residues of circulating variants) against the recombinant RBD proteins of SARS-CoV-2 WT (grey), Omicron BA.2 (blue) and KP.3.1 (red).

**Supplementary Fig. 12.**
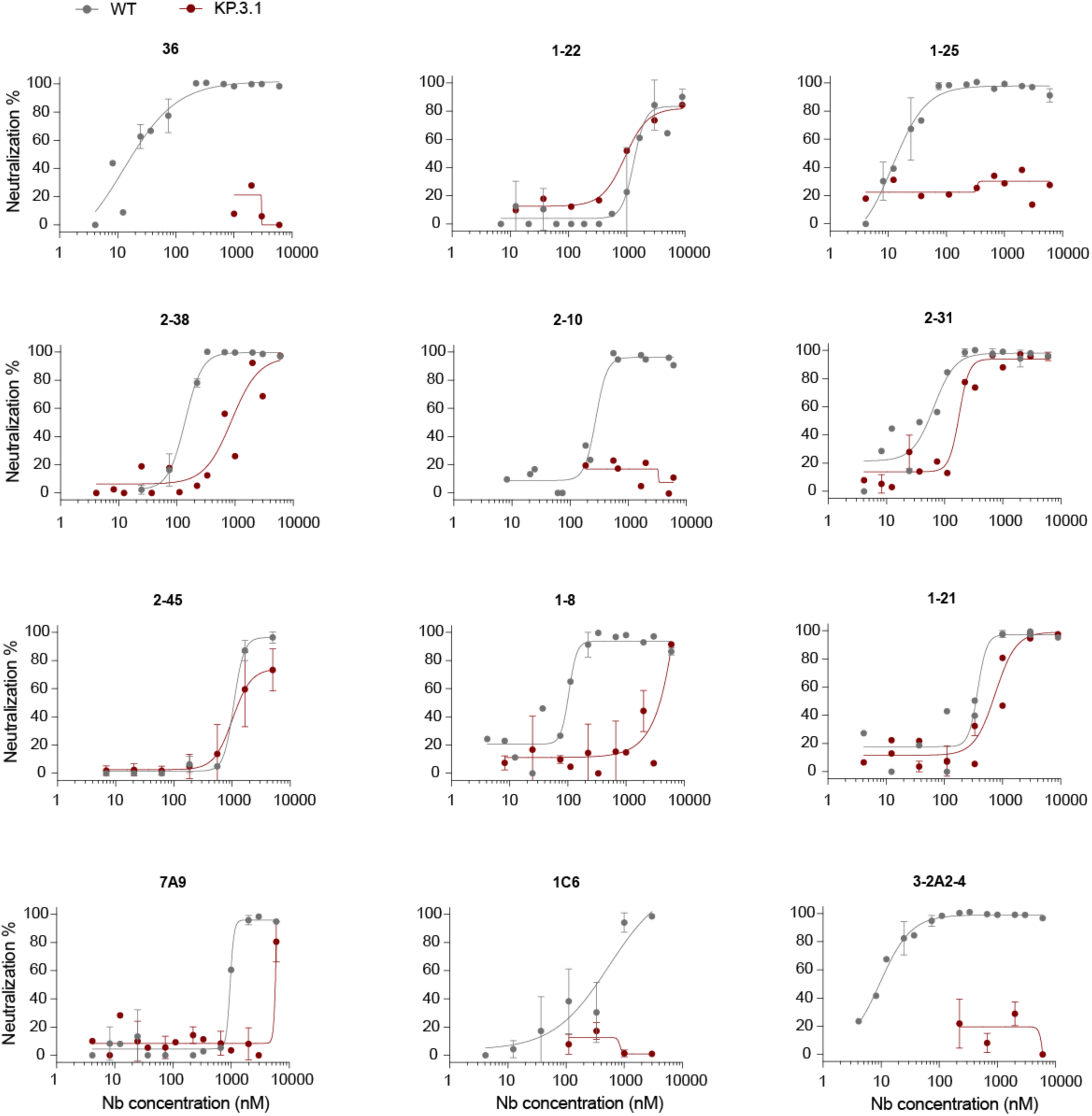
Microneutralization assay of Nbs against the authentic SARS-CoV-2 WT and KP.3.1 viruses: Three biological replicates were performed, and each dot is presented as average +/-standard deviation.

**Supplementary Fig. 13.**
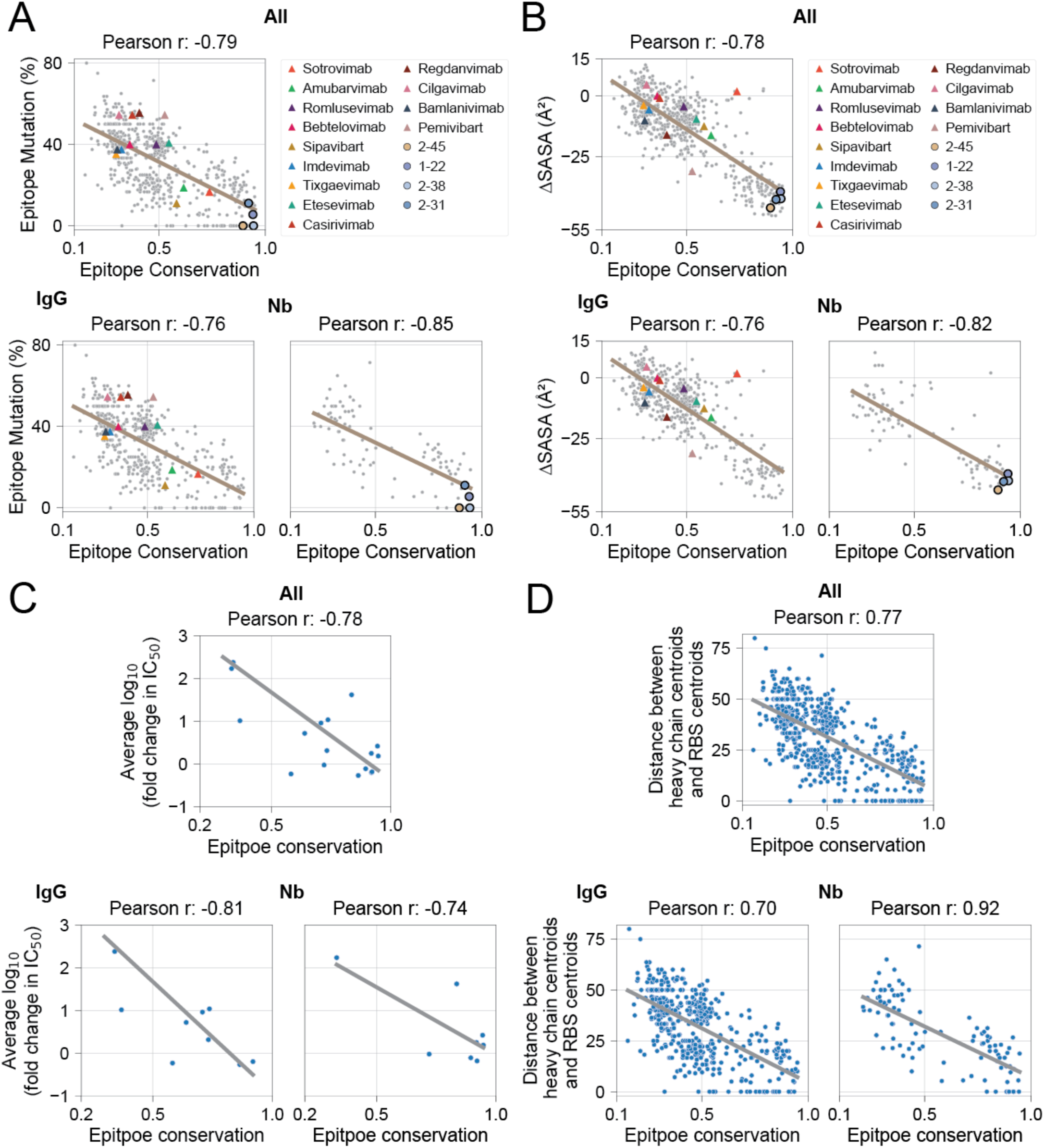
Correlation between epitope properties: (A) Upper: Correlation between epitope conservation and the percentage of epitope mutations for IgGs and Nbs. Clinical antibodies are shown as triangles, while Nbs neutralizing KP.3.1 are shown as circles. (B) Correlation between epitope conservation and epitope accessibility for IgGs and Nbs. Clinical antibodies are shown as triangles, while Nbs neutralizing KP.3.1 are shown as circles. (C) Correlation between epitope conservation and average log_10_ fold change in IC_50_ for IgGs and Nbs. (D) Correlation between epitope conservation and the distance between heavy chain centroids and RBS centroids for IgGs and Nbs.

**Supplementary Fig. 14.**
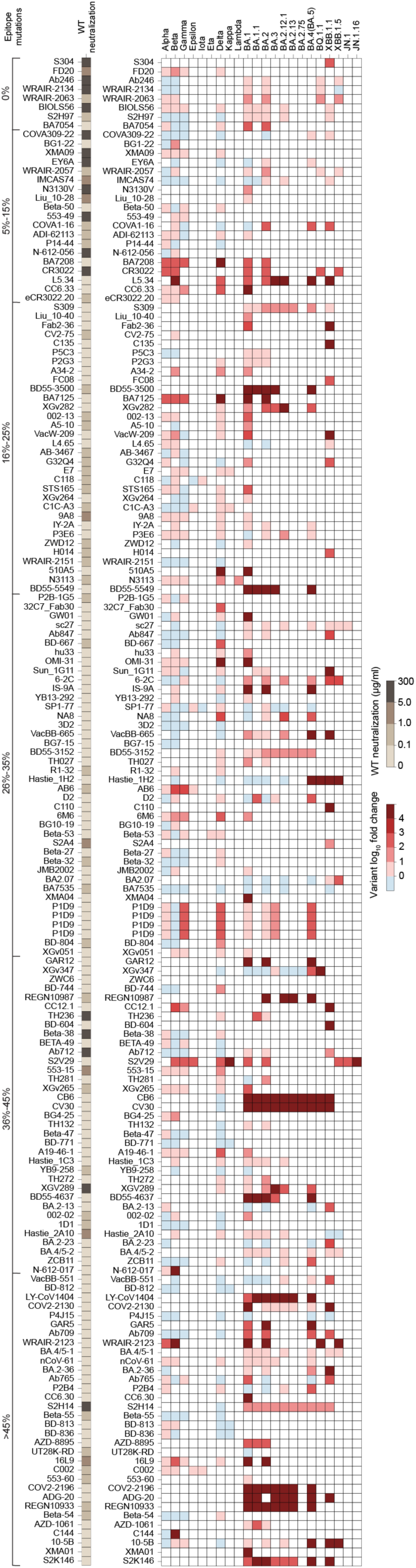
Summary of literature documents on the 174 published antibody neutralization potencies against SARS-CoV-2 and its circulating variants. Heatmap (left) of WT neutralization potency (IC_50_s) is shown in a brown gradient. Heatmap (right) displays the log₁₀ neutralization fold change against variants. Antibodies are sorted by the percentage of epitope mutations present in circulating variants. The majority of neutralization experiments were conducted using pseudovirus systems, with a few performed using authentic virus.

## Methods

### Data availability

All datasets, including structural annotations, epitope classification, and variant-associated mutation mappings, along with analysis scripts, computational tools, and interactive visualization platforms, will be made openly available to the research community.

### Antibody Structure Dataset

We collected 8556 antibody structures in PDB, as listed by SAbDab (release: 06/04/2024), including 1664 unbound and 6892 bound structures. Among bound structures, 5,742 are paired with proteins, 721 with peptides, 243 with haptens, 136 with carbohydrates, and 50 with nucleic acids. We further extracted the SARS-Cov-2 specific unique antibody structures based on PDB IDs listed in CoV-AbDab (*78*) (release: 06/04/2024), we also manually checked and added those that are missed by the Cov-AbDab leading to a total 705 bound structures for each unique antibody. In the comparison between RBD-targeting IgGs and Nbs, only naturally occurring antibodies were considered, while synthetic antibodies were excluded from the analysis based on SAbDab labels. We selected the complex structures in the highest resolution if any antibody is resolved in multiple structures. We manually renumbered the RBD domain in the complex structures to align with the numbering in the reference RBD structure (PDB: 6M0J) for later analysis by ChimeraX (*85*).

### Antigen Taxonomy Classification

To retrieve antigen taxonomy information, we obtained the InterPro (IPR) Protein Family Classification code for each protein in the structure using the PDB ID. We then obtained the taxonomy ID from the National Library of Medicine using IPR codes. The taxonomy classification was then assigned based on the ID using NCBI API.

### SARS-CoV-2 Infection Dataset

We obtained the dataset of cumulative SARS-CoV-2 infections in the U.S. from the Centers for Disease Control and Prevention (CDC) (release: 08/31/2024).

### General Structure Schematics

Superpositions, surface representations, and cartoon representations of structures were visualized by ChimeraX.

### Antibody Binding Domain and Protein

Using the SARS-CoV-2 antibody dataset described above, we screened the antigen name column to identify the specific SARS-CoV-2 protein or domain targeted by each antibody. If the antigen name does not explicitly indicate the target protein or domain, we manually reviewed the structure for verification. An antibody was defined as binding to the RBD if at least 50% of its epitope falls within the RBD region (residues 333–527).

### Identification of RBD Surface Residues

The degree of burial for a residue is quantified by measuring the depth of the side chain below the protein surface using the ResidueDepth module in BioPython. The depth of the residue side chain is calculated by average distance to surface for all atoms. The protein surface was generated by software MSMS (*79*). A cutoff of 3.03 Å defining the state of burial or exposure to solvent was used (*80*).

### Definition of Wild Type RBD Targeting Antibody

We defined an antibody as targeting the wild type (WT) RBD if it is either resolved with the WT RBD structure or its epitope does not contain any mutations associated with circulating variants.

### Identification of interface residues from antibody-RBD complex structures

An RBD residue and an antibody residue were defined in contact if the distance between any pair of their atoms was less than a threshold of 4 Å using python package ProDy (*81*). For residue-level contacts, we consider only unique residue-residue pairs as contact pairs. For atom-level contacts, we allow multiple atomic contacts between the same residue pair. The number of atomic contacts provides an indication of the strength of interaction between residues. The percentage of pairwise atomic contacts per antibody is defined as the proportion of inter-atomic contacts between specific residue pairs relative to the total number of atomic contacts in that antibody. The average pairwise atomic contact percentage is then calculated by averaging these proportions across all antibodies.

### Antibody Classification

Antibodies are classified based on the 4 epitope groups (*35*). We calculate the extent of overlap between epitope residue including VH and VL (if available) for each antibody and residue in these groups, assigning each antibody to the class with the highest percentage of residue overlap.

### RBD sequence conservation across sarbecoviruses

To calculate the conservation, 18 diverse sarbecovirus RBD sequences including WIV4, SARS-CoV, Rs7327, SHC014, RaTG13, pang17, Yun11, Rs4081, Rf1, Rf4092, YN2013, ZC45, HKU3-1, Shanxi2011, Rp3, BM4831, BtKY72, were aligned to SARS-CoV-2 RBD sequence. After alignment, each SARS-CoV-2 RBD residue was compared to corresponding residue in other RBD sequences. An identical residue to that in SARS-CoV-2 is considered a match. The conservation for each RBD residue is calculated by the number of matches over 18. To estimate average epitope conservation, we defined epitope residues as residues with at least one atom within 4Å from the antibody atom and averaged conservation over epitope residues.

### Viral fitness score of epitope residues

The viral fitness score was obtained from (*38*) by evaluating the mutational effects on RBD expression level. The fitness score of each RBD residue is the averaged value of mutating to any other amino acids. The viral fitness score of epitopes was calculated by averaging fitness score over epitope residues. To estimate average epitope viral fitness, we defined epitope residues as residues with at least one atom within 4Å from the antibody atom and averaged viral fitness over epitope residues.

### Epitope Surface Area

The antibody epitope residues were extracted based on the method described above. Epitope residues were defined as those with at least one atom within 4 Å of an antibody atom. The solvent-accessible surface area (SASA) of the molecules was calculated using python package FreeSASA (*82*), and the SASA of the epitope residues was summed to determine the epitope size.

### Epitope accessibility in context of RBD dynamics on Spike

RBD surface residues have different degrees of solvent accessibility between RBD_down_ and RBD_up_ conformations, we calculate the difference of solvent accessible surface area (ΔSASA) for RBD residues between these two conformational states to quantify the RBD surface accessibility. Specifically, solvent-accessible surface area (SASA) was calculated using the python package FreeSASA for the RBD in both the “down” conformation (PDB: 6XR8) and the “up” conformation (PDB: 7X7N). Epitope residues were defined as RBD residues having at least one atom within 4 Å of any antibody atom in complex structures. ΔSASA was quantified as the difference in the total SASA of epitope residues between the RBD-down and RBD-up conformations. ΔSASA = SASA_down_ − SASA_up_, where a negative value indicates decreased surface exposure in the down conformation.

### Identification of CDR/FR sequences in antibody/nanobody

Antibody (VH/VL) or nanobody sequences were numbered by the python package ANARCI (*83*) under the IMGT numbering scheme, the CDR/FR sequences were identified based on IMGT definition.

### Circulating variant mutations

Mutational data and residues for SARS-CoV-2 circulating variants were obtained from Outbreak.info (*84*).

### Percentage of epitope mutations

Interface residues are defined as described above. The percentage of epitope mutations was calculated by comparing the number of epitope mutations in the RBDs of the circulating variants to WT and then divided by the total number of epitope residues. For antibody–SARS-CoV-2 RBD complex structures that are not solved with WT RBD, comparisons were made between the RBD of variants and the RBD in the complex instead.

### Superposition of CDR3 loops within epitope sub-cluster

Complexes of antibody CDR3 loops and RBD were extracted from original structures using in house python scripts. In brief, anchors of CDR3 loops are first defined to be N-terminal of FR3 (IMGT numbered residue 102-104, e.g. YYC) and N-terminal of FR4 (IMGT numbered residue 118-121, e.g. WGQG). Structures of CDR3 loop and its anchor along with the RBD are then extracted. To visualize the CDR3 loop conformations within the same epitope subcluster, extracted structures are superimposed with reference to RBD. To compare the CDR3 loop conformation, CDR3 loops belonging to the same subcluster are superimposed with reference to the anchor regions.

### Quantification of CDR3 loop sequence similarity

The sequence identity between two sequence is calculated by MMSeqs2 (*88*). The sequence similarity within the same group or between different groups is then calculated as follows:

#### a) Intra-group comparison

Each sequence is aligned with all other sequences within the same group,then the average of sequence identities from alignments is calculated. The intra-group sequence similarity is represented as a distribution of average sequence identities.

#### a) Inter-group comparison

The comparison for inter-group sequence similarity is largely identical to intra-group comparison, however instead of aligning sequences from the same group, each sequence is aligned to all sequences from another group.

### Published IC_50_ Data for WT and Delta Variant Neutralization

IC50 data for antibody neutralization against the SARS-CoV-2 Delta variant and WT were extracted from published experimental results reported in (*88–95*).

### Epitope similarity in analyzing heavy chain CDR subclusters

To quantify epitope similarity between two antibodies, we computed the intersection-over-union (IoU) of their respective VH epitope residues. Specifically, let E_1_ and E_2_ denote the sets of epitope residues recognized by the two antibodies. The epitope similarity is defined as:

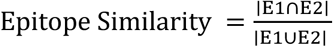

This metric ranges from 0 (completely distinct epitopes) to 1 (identical epitopes) and provides a normalized measure of epitope similarity between antibody pairs. We used 0.4 epitope similarity as a threshold for subclusters in Fig. 3.

### Source of Clinical Antibody Regulatory Status

Approval and revocation dates of clinical antibodies were obtained from official fact sheets and reports provided by the U.S. Food and Drug Administration and the European Medicines Agency.

### Preparation of Starting Structures for MD simulations

Starting structures for the wild-type RBD were derived from the corresponding PDB structures using the comparative modelling software MODELLER (**Table S5**) (*96*). This approach ensured consistency across structures resolved by various experimental techniques and software. When the complex structure was resolved using x-ray crystallography, it was used directly as a template for MODELLER. In cases where cryo-EM structures had poorly resolved RBD regions, we substituted the RBD with that from a high-resolution x-ray structure (PDB 7JVB), creating a refined template for MODELLER. For all other RBD variants, the starting structures were computed by MODELLER starting from the corresponding WT RBD complex as a template. Additionally, for one nanobody complexes (Nb2-57) with poorly resolved CDR loops, AlphaFold3 was used to remodel the nanobody which was then superimposed onto the original nanobody structure in the PDB file.

### MD Simulations

The MD simulations were performed with GROMACS 2024 software (*97*) using the CHARMM36m force field (*98*). Each of the complexes was solvated in transferable intermolecular potential with 3 points (TIP3P) water molecules, and ions were added to equalize the total system charge. The steepest descent algorithm was used for initial energy minimization until the system converged at Fmax < 1000 kJ/(mol nm). Then, water and ions were allowed to equilibrate around the protein in a two-step equilibration process. The first part of the equilibration was at a constant number of particles, volume, and temperature (NVT). The second part of the equilibration was at a constant number of particles, pressure, and temperature (NPT). For both of the MD equilibration parts, positional restraints of k = 1000 kJ/(mol nm2) were applied to the heavy atoms of the protein, and the system was allowed to equilibrate at a reference temperature of 300 K, or reference pressure of 1 bar for 100 ps at a time step of 2 fs. Following the equilibration, the production simulation duration was 100 nanoseconds with 2 fs time intervals. Altogether 10,000 frames were saved for the analysis at intervals of 10 ps.

### Metrics and Scoring

To assess the complex-wide structural change induced by the simulation, RMSD values between all frames and the initial structure were computed. In examining the interaction between the viral RBD and the nanobodies at the interface, multiple techniques were employed, which were focused on each frame independently. Statistically optimized atomic potentials (SOAP) scoring (*99*) was used to measure potential energy at the interface based on atomic distance. Interface contact area and interface energy (VoroIF-jury) (*100*) were used as well, which are two measures computed by the Voronota tool whose technique relies on Voronoi tessellation of protein structures. The former is a geometric measure which computes the surface area of contact between pairs of atoms at the interface, as given by the tessellation, where each atom in a pair belongs to a different chain. The latter is a pseudo-energetic approximation derived from the tessellation and accounts for pairs of atoms in contact at the interface similarly to the surface contact area.

### Nanobody Synthesis and Purification

The monomeric Nb genes were codon-optimized and synthesized from Synbio Technologies. All the Nb DNA sequences were cloned into a pET-21b(+) vector using EcoRI and HindIII restriction sites. Nb DNA constructs were transformed into Shuffle T7 cells (NEB) and plated on Agar with 50µg/mL ampicillin at 37C overnight. Single clones were picked and cultured in an LB broth to reach an O.D. of 0.6–0.8 before IPTG (0.5-1mM) induction at 16C overnight. Cells were then harvested, sonicated, and lysed on ice with a lysis buffer (1xPBS pH7.4, 150 mM NaCl, 0.2% TX-100 with protease inhibitor). After cell lysis, protein extracts were collected by centrifugation at 21,000 x g for 10 mins and the his-tagged Nbs were purified by the Cobalt resin (Thermo) and natively eluted with a buffer containing 150 mM imidazole buffer. Eluted Nbs were subsequently dialyzed to 1x DPBS, pH 7.4 using the Amicon ultra centrifugal units (3kda cut-off).

### Production of Recombinant SARS-CoV-2 RBD Proteins

Plasmid of the WT RBD of SARS-CoV-2 (GenBank MN985325.1; S protein residues 319–539) was synthesized from Synbio Technologies. To express the protein, Expi293F cells were transiently transfected with the plasmid using the ExpiFectamine 293 kit. After 20 hrs of transfection, enhancers were added to further boost protein expression. Cell culture was harvested 3–4 days after transfection and the supernatant was collected by high-speed centrifugation at 21,000xg for 30 min. The secreted proteins in the supernatant were purified using His-Cobalt resin. Eluted proteins were further purified by size-exclusion chromatography using a Superdex 75 Increase column (Cytiva) in a buffer composed of 20 mM HEPEs pH 7.2 and 150 mM NaCl. SARS-CoV-2 RBD variants BA.2 and KP.3 were obtained from the Acro Biosystems and Sino Biological.

### ELISA (Enzyme-Linked Immunosorbent Assay)

Indirect ELISA was carried out to evaluate the relative affinities of the Nbs against different RBDs. A 96-well ELISA plate (R&D system) was coated with the RBD protein at an amount of 5 ng per well in a coating buffer (15 mM sodium carbonate, 35 mM sodium bicarbonate, pH9.6) overnight at 4C, with subsequent blockage with a blocking buffer (1xPBS pH 7.4, v/v0.05% Tween20, 5% milk) at room temperature for 2 hours. Nbs were serially 5-fold diluted from 1µM to 12.8pM in the blocking buffer. The dilutions were incubated for 2 hours at room temperature. HRP-conjugated secondary antibody cocktail against the VHH (Genscript) was diluted at 1:5,000 in the blocking buffer and incubated with each well for 1 hour at room temperature. Three washes with 1xPBST (1xPBS pH7.4, v/v0.05%Tween20) were carried out to remove non-specific absorbances between each incubation. After the final wash, the samples were further incubated in the dark with freshly prepared w3,30,5,50-Tetramethylbenzidine (TMB) substrate for 10 mins at room temperature to develop the signals. After the STOP solution (R&Dsystem), the plates were read at multiple wavelengths (450nm and 550nm) on a plate reader (Thermo Fisher). The raw data were processed by Prism 9 (GraphPad) to fit into a 4PL curve and to calculate log IC_50_.

### Cell Lines

VeroE6 cell line constitutively expressing full-length human TMPRSS2 (Vero-TMPRSS2) was purchased from BPS Bioscience (78081) and maintained in Dulbecco’s modified Eagle’s medium (DMEM) with glucose, L-glutamine, and sodium pyruvate (Corning, 10-017-CV) supplemented with 10% fetal bovine serum (FBS, Peak Serum), non-essential amino acids (Corning, 25-025-CI), penicillin (100 UI/mL) and streptomycin (100 UI/mL) (Corning, 30-002-CI), and puromycin (3 ug/mL) (InvivoGen, ant-pr-1) (cDMEM). Cells were grown at 37°C in 5% CO_2_ and supplemented with normocin (100 ug/mL) (InvivoGen, ant-nr-1) to prevent mycoplasma infection.

### Viruses

SARS-CoV-2 ancestral WA1^614G^ and Omicron KP.3.1 variants used in this study were originally isolated from nasopharyngeal swabs of COVID-19 infected individuals as previously described (*101*) and kindly provided by Dr. Viviana Simon. Specimens were collected as part of the routine SARS-CoV-2 surveillance conducted by the Mount Sinai Pathogen Surveillance programme (IRB approved, HS#13-00981). To generate viral stocks, Vero-TMPRSS2 cells were infected at a MOI of 0.01 and maintained in infection media (DMEM with glucose, L-glutamine, and sodium pyruvate supplemented with 2% FBS, non-essential amino acids, HEPES, penicillin (100 UI/mL) and streptomycin (100 UI/mL)) at 37°C in 5% CO_2_. Cells were monitored for cytopathic effect, and viral supernatants were collected at day 3 post-infection for WA1^614G^ and at day 4 post-infection in the case of Omicron KP3.1 subvariant. Viral supernatants were clarified of cell debris by spin down and aliquots were stored at −80°C. All viral stocks were sequence-confirmed by next-generation sequencing. Experiments with authentic SARS-CoV-2 viruses were performed in the biosafety level 3 (BSL-3) facility following Icahn School of Medicine biosafety guidelines.

### Microneutralization Assay

Microneutralization assays were performed as previously described(*61, 62*) with modifications. Briefly, Vero-TMPRSS2 cells were seeded at a density of 20,000 cells per well in a 96-well cell culture plate in cDMEM. The following day, IgGs and Nbs (dilution of 1:10) were serially diluted three-fold in 1x minimum essential medium (MEM) with 2% FBS in a final volume of 200 µl. Next, SARS-CoV-2 variants were diluted to a concentration of 100 Tissue Culture Infectious Dose 50 percent (TCID_50_) in 1× MEM. 80 µl of serum dilution and 80 µl of virus dilution were added to a new 96-well plate and incubated for 1 h at 37 °C. Then, cDMEM was removed from Vero-TMPRSS2 cells and 120 µl of virus-serum mixture was added to the cells. The cells were incubated at 37 °C for 1 h. After 1 hour incubation, virus-serum mixture was removed from the cells and 100 µl of corresponding serum dilutions and 100 µl of 1xMEM with 2% FBS was added to the cells. Then, cells were incubated for 24 h and fixed with 10% paraformaldehyde (Polysciences) for 24h at 4 °C. Following fixation, cells were washed with phosphate-buffered saline (Corning) with Tween-20 (Fisher) (PBST) and permeabilized with 0.1% Triton X-100 (Fisher) for 15 min at room temperature. The cells were washed three times with PBST and blocked with 3% milk in PBST for 1 h at room temperature. Then, the cells were incubated with mouse antibody 1C7 (anti-SARS nucleocapsid antibody, kindly provided by Dr. Moran) at a dilution of 1:1000 in 1% milk in PBST and incubated for 1 h at room temperature. The cells were washed three times with PBST. Then, cells were incubated with goat anti-mouse IgG-HRP (Abcam, Cat. ab6823) at a dilution of 1:10,000 in 1% milk in PBST and incubated for 1 h at room temperature. The cells were washed three times with PBST and TMBE Elisa peroxidase substrate (Rockland) was added. After 15 min incubation, sulfuric acid 4.0 N (Fisher) was added to stop the reaction, and the readout was done using a Synergy H1 plate reader (BioTek) at an OD450.

## Acknowledgments

We thank Dr. Randy Albrecht for support with the BSL3 Conventional Biocontainment Facility (CBF) and procedures at the ISMMS. We are indebted to the efforts of the Mount Sinai Pathogen Surveillance Program (PSP) for the collection and sequencing of clinical specimens used to derive SARS-CoV-2 isolates. We also thank Shi lab members for their valuable input and discussion. Funding: This work is supported by NIH grant R01 AI163011 (Y.S. and D.S.). This work is also partly supported by CRIPT (Center for Research on Influenza Pathogenesis and Transmission), an NIAID funded Center of Excellence for Influenza Research and response (CEIRR, contract # 75N93021C00014), and by NIAID grant U19AI135972 (A.G.-S.). The Conventional Biocontainment Facility (CBF) is a NIHBSL3/BSL3 facility that is part of the BSL-3 Biocontainment CoRE at ISMMS. This core is supported by subsidies from the ISMMS Dean’s Office and by investigator support through a cost recovery mechanism. Research reported in this publication was supported by NIAID Award Number G20AI174733 (R.A. Albrecht). The content is solely the responsibility of the authors and does not necessarily represent the official views of the National Institutes of Health.

## Author contributions

Y.S. conceived the study and all analyses. Z.F. and Z.S. developed the scripts supporting data analysis. Z.F. performed the majority of data analysis and Fig. generation. A.W. conducted MD simulations. Y.X. and A.E. carried out the biochemical and neutralization experiments. Y.S., A.G.-S., and D.S. supervised the overall study. Y.S. and Z.F. drafted the manuscript with input from all authors.

